# *De novo* emergence of adaptive membrane proteins from thymine-rich intergenic sequences

**DOI:** 10.1101/621532

**Authors:** Nikolaos Vakirlis, Omer Acar, Brian Hsu, Nelson Castilho Coelho, S. Branden Van Oss, Aaron Wacholder, Kate Medetgul-Ernar, John Iannotta, Aoife McLysaght, Carlos J. Camacho, Allyson F. O’Donnell, Trey Ideker, Anne-Ruxandra Carvunis

## Abstract

Recent evidence demonstrates that novel protein-coding genes can arise *de novo* from intergenic loci. This evolutionary innovation is thought to be facilitated by the pervasive translation of intergenic transcripts, which exposes a reservoir of variable polypeptides to natural selection. Do intergenic translation events yield polypeptides with useful biochemical capacities? The answer to this question remains controversial. Here, we systematically characterized how *de novo* emerging coding sequences impact fitness. In budding yeast, overexpression of these sequences was enriched in beneficial effects, while their disruption was generally inconsequential. We found that beneficial emerging sequences have a strong tendency to encode putative transmembrane proteins, which appears to stem from a cryptic propensity for transmembrane signals throughout thymine-rich intergenic regions of the genome. These findings suggest that novel genes with useful biochemical capacities, such as transmembrane domains, tend to evolve *de novo* within intergenic loci that already harbored a blueprint for these capacities.

---

The molecular mechanisms and dynamics of *de novo* gene birth are poorly understood^1^. It is particularly unclear how non-genic sequences could spontaneously encode proteins with specific and useful biochemical capacities. To resolve this paradox, it has been proposed that pervasive translation of non-genic transcripts can expose genetic variation, in the form of novel polypeptides, to natural selection, thereby purging toxic sequences and providing adaptive potential to the organism^2, 3^. The genomic sequences encoding these novel polypeptides have been called “proto-genes”, to denote that they correspond to a distinct class of genetic elements that are intermediates between non-genic sequences and established genes^3^. In agreement, several studies reported that *de novo* emerging coding sequences tend to display lengths, transcript architectures, transcription levels, strength of purifying selection, sequence compositions, structural features and integration in cellular networks that are intermediate between those observed in non-genic sequences and those observed in established genes^3–8^. Furthermore, pervasive translation of non-genic sequences has been observed repeatedly by ribosome profiling and proteo-genomics^3, 9–12^, and studies have shown that random sequence libraries harbor bioactive effects^13–17^. Nonetheless, whether and how native proto-genes carry adaptive potential remains unknown.

We sought to formalize the predictions of adaptive proto-gene evolution. We define adaptive potential as the capacity to increase fitness by means of evolutionary change. While any sequence may in theory carry adaptive potential, changes in established genes are typically constrained by preexisting selected effects – the specific physiological processes mediated by the gene products that lead to their evolutionary conservation^18^. In contrast, emerging proto-genes are expected to mostly lack such selected effects, leaving them more readily accessible to adaptive change and innovation^2, 3^. We reasoned that the initial adaptive potential would give way, as proto-genes mature and the adaptive changes engender novel selected effects, in turn reducing the possibility of future change. This reasoning is akin to Sartre’s “existence precedes essence” dictum^19^, and predicts that proto-genes are enriched in adaptive potential and depleted in selected effects relative to established genes (Fig. 1A).

**Fig. 1.**
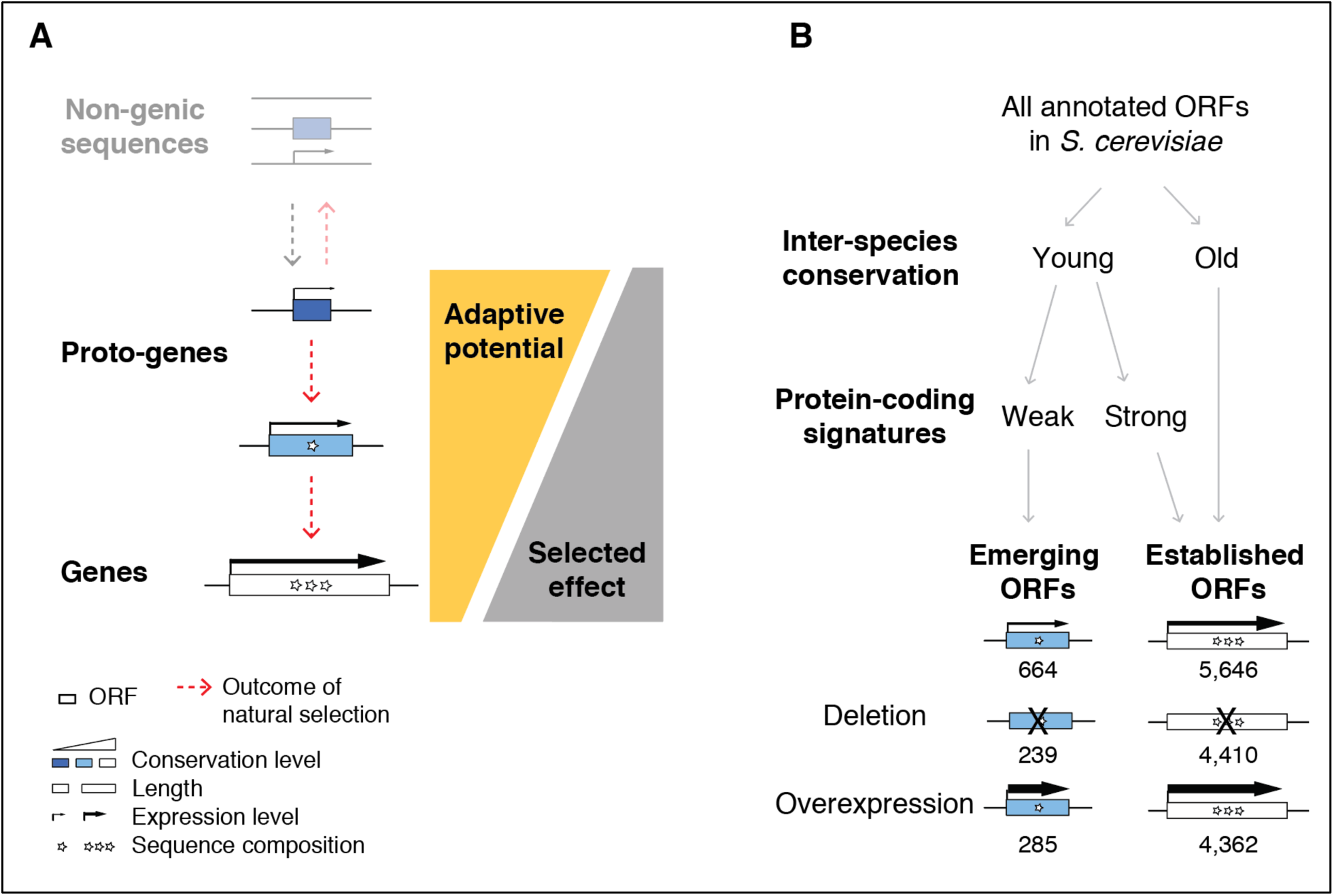
The adaptive potential prediction: theory and empirical testing. **A.** <u>Theoretical model</u>. The evolution from non-genic sequences to proto-genes to genes is represented as in ref^3^; the transition from non-genic sequences to proto-genes is mediated by the act of translation; the transition from proto-genes to genes occurs along a continuum (left). Proto-genes are predicted to provide adaptive potential to the organism by exposing natural variation to selection, while being depleted in selected effects (right). **B.** <u>Operational classification of emerging and established ORFs</u>. We confront the theoretical model by systematically assessing how emerging ORFs impact fitness in budding yeast. Emerging ORFs are young (no detectable homologues outside of the *sensu stricto* genus and no conserved syntenic homologue in *S. kudryiavzevii* and *S. bayanus*) and do not display strong evidence that they encode a useful protein product under intraspecific purifying selection (**Methods**). Empirical testing of the theoretical prediction (**A**) involves experimentally measuring the fitness of deletion and overexpression alleles for both classes of ORFs. The numbers of emerging and established ORFs subjected to each analysis are indicated.

In what follows, we confronted these theoretical predictions with systematic measurements of how disruption and overexpression of open reading frames (ORFs) impact fitness in budding yeast as a function of the evolutionary emergence status of the ORFs. We classified annotated *S. cerevisiae* ORFs into emerging ORFs and established ORFs (Fig. 1B; **Data S1**). The established ORFs group contained ancestral ORFs with high interspecific conservation levels, as well as evolutionarily young ORFs that encode useful proteins under intraspecific purifying selection. The emerging ORFs group contained evolutionarily young ORFs whose recent evolution was determined by phylostratigraphy and syntenic alignments, and which lacked strong evidence of encoding a useful protein product (**Methods**). As expected, emerging ORFs tend to be short and weakly transcribed relative to established ORFs (Mann-Whitney U test *P*<2.2×10^−16^ in both cases). Most emerging ORFs (>95%) are annotated as Dubious or Uncharacterized by domain experts (**Methods**), as it is currently unclear whether they correspond to spurious non-genic ORFs or to young genes whose physiological implications remain to be discovered.

## Selected effects

We compared estimated fitness costs of disrupting emerging and established ORFs. To this aim, we first examined fitness estimates generated from a large collection of systematic deletion and hypomorphic alleles^20^. After removing ORFs with overlapping genomic locations, we obtained fitness estimates for 239 emerging and 4,410 established ORFs (Fig. 1B). Fitness estimates were markedly higher when comparing deletions of emerging ORFs to deletions of established ORFs (Mann-Whitney U test *P*=1.5×10^−17^). For example, only 8% of emerging ORFs were associated with even a small fitness cost (n=19; fitness estimate<0.9), relative to 29% of established ORFs (n=1,290; Fisher’s exact test *P*<3.6 ×10^−15^) (Fig. 2A). The low fitness cost of disrupting emerging ORFs in laboratory conditions was consistent with their low native expression level (Fisher’s exact test *P* = 0.09), and more pronounced than expected compared to established ORFs of matched length distribution (Fisher’s exact test *P* = 3.6×10^−15^) (Extended Data Fig. 1A).

**Fig. 2.**
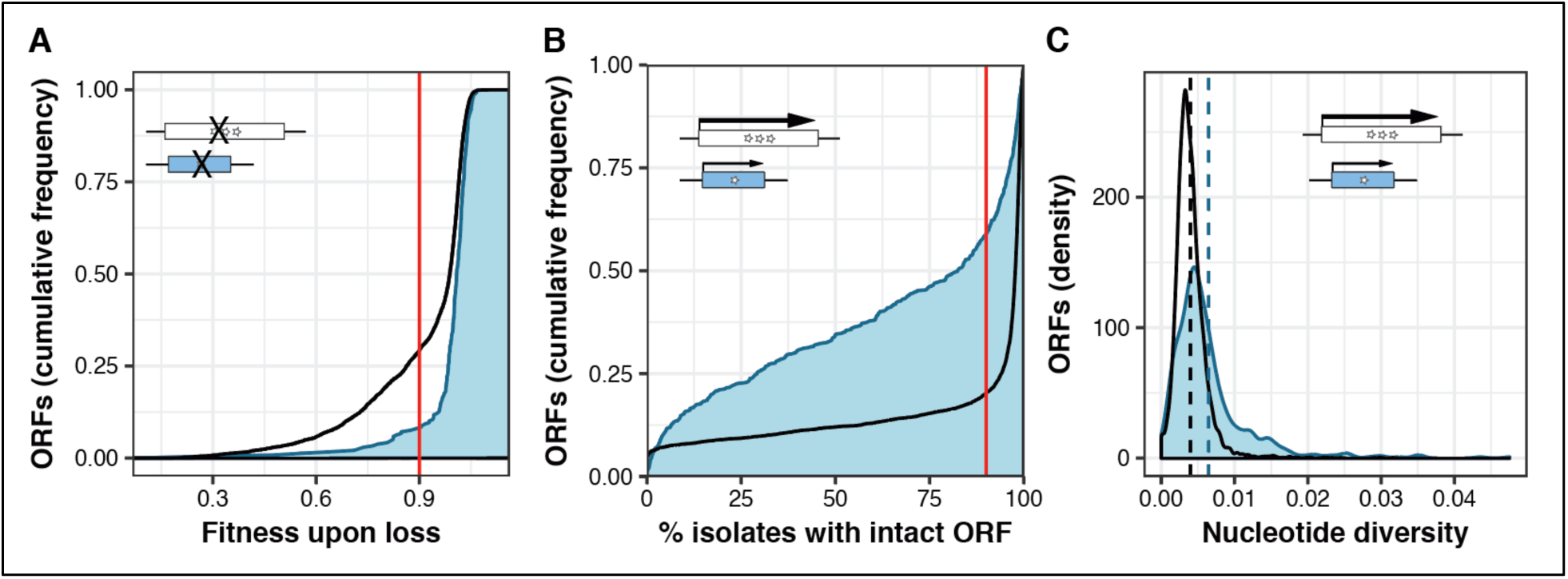
Disruption of emerging ORFs is generally inconsequential for fitness. **A.** <u>Deletion of emerging ORFs incurs lesser fitness costs than deletion of established ORFs.</u> Empirical cumulative distribution function for emerging (blue) and established (black) ORFs; fitness of deletion strains in rich media (YPD) at 30°C as estimated by ref^20^, averaged over multiple alleles per ORF when applicable. Vertical red line illustrates the fraction of ORFs of each group with fitness less than 0.9. **B.** <u>ORF structures are more variable for emerging than established ORFs across 1,011 *S. cerevisiae* isolates.</u> Empirical cumulative distribution function for emerging (blue) and established (black) ORFs; ORF structure defined as intact in a pairwise alignment if the positions of the start codon and stop codons are maintained, the frame is maintained, and intermediate stop codons are absent. Vertical red line illustrates the fraction of ORFs of each group found intact in less than 90% of isolates. **C.** <u>Emerging ORFs display higher nucleotide diversity than established ORFs across *S. cerevisiae* isolates</u>. Density distributions for emerging (blue) and established (black) ORFs; nucleotide diversity estimated over multiple alignments lacking unknown base calls exclusively. Vertical dashed lines represent group means.

To investigate how the disruption of emerging ORFs impacts fitness in natural conditions, we analyzed intraspecific sequence variation across 1,011 *S. cerevisiae* isolates^21^. Counting the number of isolates in which the ORF structures (defined as start, stop and frame without considering sequence similarity) were intact in each group, we found ORF structures to be markedly more variable across isolates for emerging than established ORFs (Fig. 2B), including established ORFs with matched length and expression level distributions (Fisher’s exact test *P* < 2.1×10^−15^ in both cases) (Extended Data Fig. 1B). For example, while 80% of established ORFs were intact in more than 90% of isolates, indicating that they are fixed within the species, this was only the case for 41% of emerging ORFs. Furthermore, estimates of within-species nucleotide diversity were markedly higher for emerging than established ORFs (Mann-Whitney U test *P*=6.4×10^−46^) (Fig. 2C). Altogether, our results confirmed that disrupting emerging ORFs is generally inconsequential for survival in both laboratory and natural settings, as expected for loci that lack evidence of encoding a useful protein product. These findings argue against the notion that emerging ORFs might correspond primarily to young established genes whose physiological implications remain to be discovered.

## Adaptive potential

Across kingdoms, one type of evolutionary change that typically accompanies *de novo* gene birth is an increase in expression level^22^. It follows that, according to our prediction (Fig. 1A), increasing the expression level of emerging ORFs should increase the organism’s fitness more frequently than when the same perturbation is imposed on established ORFs (whose expression levels have presumably been optimized by natural selection). Alternatively, if emerging ORFs mostly correspond to spurious non-genic ORFs with no role in *de novo* gene birth, increasing their expression level should generally be neutral or toxic, and not provide fitness benefits. Systematic overexpression screens have been shown to recapitulate the outcomes laboratory evolution experiments^23^. We thus developed a dedicated overexpression screening strategy to identify ORFs that increased relative fitness upon increased expression, whereby colony sizes of individual overexpression strains were compared with those of hundreds of replicates of a reference strain with the same genetic background on ultra-high-density arrays (Fig. 3A; **Methods**).

**Fig. 3.**
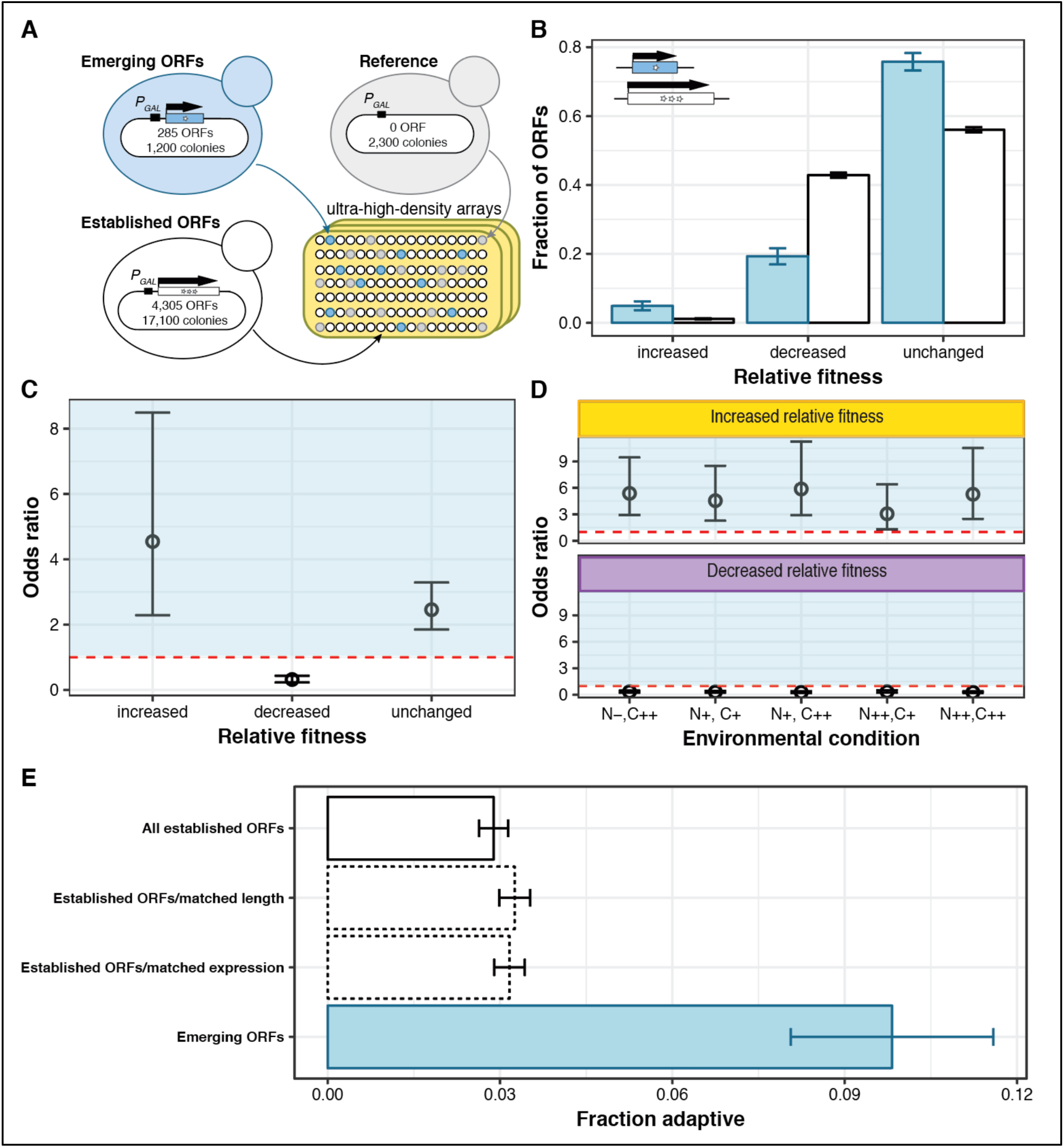
Testing the adaptive potential prediction by estimating relative fitness upon overexpression. **A.** <u>Strategy to screen for relative fitness effects of overexpression strains.</u> Yeast strains overexpressing emerging and established ORFs, and reference strains, are arrayed at ultra-high-density on plates containing agar media. Fitness is estimated from the distributions of colony sizes of technical replicates. Number of colonies are rounded to the nearest hundred. See **Methods**. **B.** <u>Fraction of emerging (blue) and established (white) ORFs displaying increased, decreased and unchanged fitness effects relative to the reference.</u> Environmental condition was SC-URA+GAL+G418 media (**Table S1**). Error bars represent standard error of the proportion. **C.** <u>Emerging ORFs are 4.5 times more likely to increase relative fitness when overexpressed than established ORFs, and 3.1 times less likely to decrease relative fitness.</u> Odds ratios derived from the data shown in panel **B**. Vertical error bars represent 95% confidence intervals. Horizontal dashed line indicates odds ratio of 1. All odds ratios are significantly different from 1 (P<0.00002). **D.** <u>Emerging ORFs are consistently more likely to increase fitness and less likely to decrease fitness than established ORFs when overexpressed in five different environments.</u> “N”: poor (-), complete (+) or rich (++) supplementation of amino-acids; C: complete (+) or rich (++) supplementation of carbon sources (**Table S1**). Odds ratios represent the likelihood of emerging ORFs to increase or decrease relative fitness compared to established ORFs. Vertical error bars represent 95% confidence intervals. Horizontal dashed lines indicate odds ratio of 1. All odds ratios are significantly different from 1 (*P*<0.00002). **E.** <u>Proportion of ORFs displaying increased fitness relative to the reference in at least one of five different environments.</u> Blue: emerging ORFs; White with solid contour line: established ORFs; White with dashed contour line: established ORFs sampled with replacement according to the distribution of ORFs lengths and native RNA expression levels of emerging ORFs. While sampling shorter or less expressed established ORFs did marginally increase the proportion found adaptive, none of these factors was sufficient to explain the high proportion of emerging ORFs found adaptive. Error bars represent standard error of the proportion.

We deployed our screening strategy in a plasmid-based overexpression collection^24^ containing 285 emerging ORFs and 4,362 established ORFs (Fig. 1B), having verified that the presence of an overexpression plasmid did not lead to a detectable growth defect relative to a plasmid-free strain (Extended Data Fig. 2; **Methods**). Strains overexpressing 14 emerging and 49 established ORFs displayed increased relative fitness in complete media, representing 4.9% and 1.1% of the total number of emerging and established ORFs tested, respectively (Fig. 3B). Overall, overexpressing most ORFs did not significantly change colony sizes relative to the reference strain. Nevertheless, overexpression of emerging ORFs was 4.5 times more likely to increase relative fitness, and 3.1 times less likely to decrease relative fitness, than overexpression of established ORFs (Fig. 3C). The tendency of emerging ORFs to increase fitness when overexpressed was also observed in the context of a pooled competition in the same media (Mann-Whitney U test *P*=5.5×10^−32^) (Extended Data Fig. 3A). Emerging ORFs with increased relative fitness displayed effect sizes ranging from 7.9% to 19% (Extended Data Fig. 3B), which is remarkable since adaptive mutations resulting in a 10% fitness increase are estimated to reach 5% of the population in ∼200 generations and fix in ∼500 generations^23^. One of the beneficial emerging ORFs identified by our experiments was *MDF1* (YCL058C), one of the best-studied examples of adaptive *de novo* origination^25, 26^.

Expanding our screening strategy to five environments of varying nitrogen and carbon composition (**Table S1; Data S3**), we found that strains overexpressing emerging ORFs were consistently 3-to 6-fold more likely to increase relative fitness and 3-to 4-fold less likely to decrease relative fitness, compared to strains overexpressing established ORFs, across all environments tested (Fig. 3D). Notably, while overexpression of only 2.9% of established ORFs increased relative fitness in at least one environment (n=126), this was the case for 9.8% of emerging ORFs (n=28) (Fig. 3E, Extended Data Fig. 4). Sixty percent (17/28) of these adaptive emerging ORFs provided fitness benefits across two or more environmental conditions (empirical *P*-value <10^−5^; Extended Data Fig. 4). The strong over-representation of adaptive effects in emerging ORFs relative to established ORFs (Fisher’s exact test *P* = 1.2×10^−7^; Odds ratio: 3.7) could not be explained by their short length or low native expression levels (Fig. 3E).

**Fig. 4:**
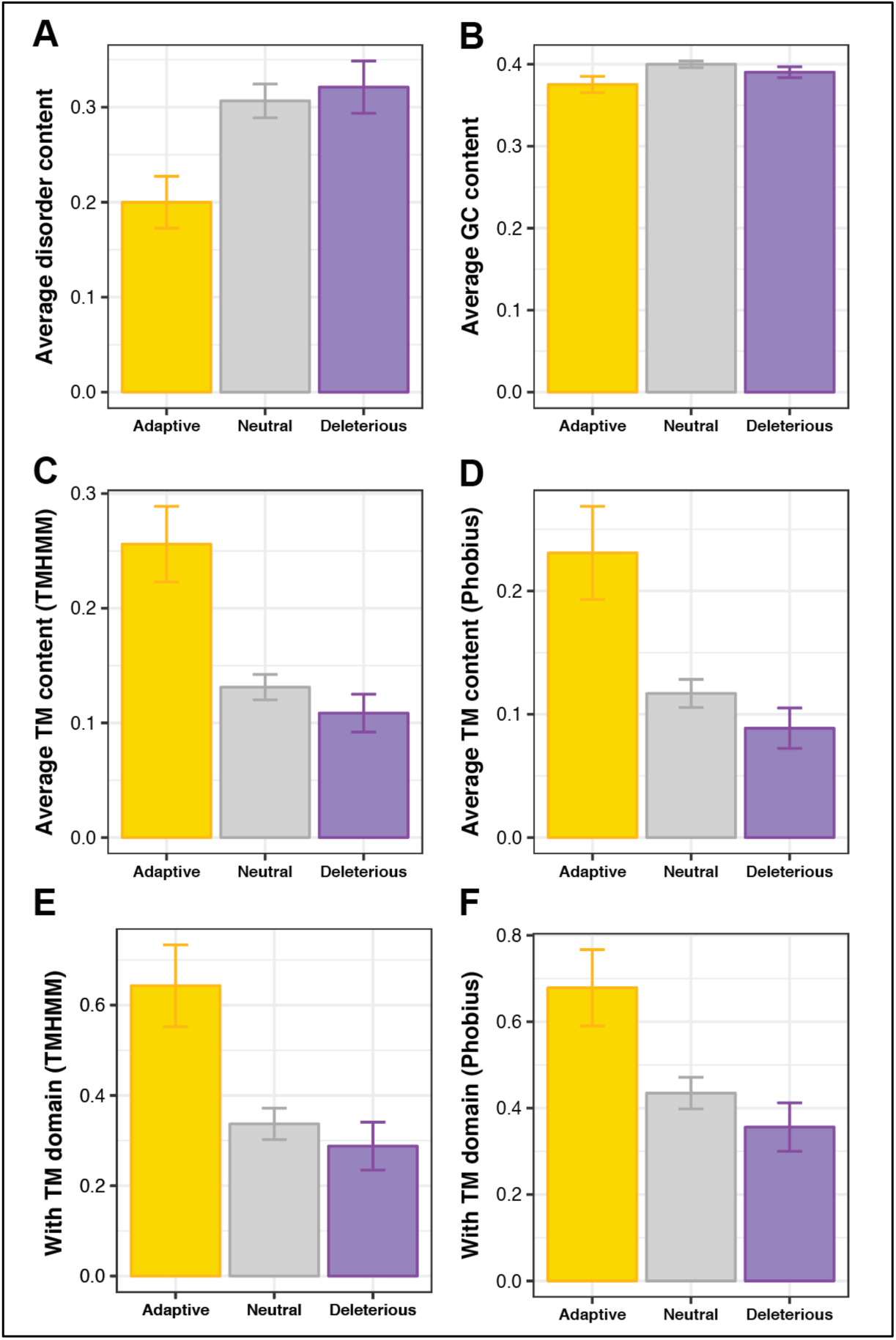
TM propensity is associated with beneficial fitness effects in emerging ORFs. **A.** <u>No association between high disorder and beneficial fitness effects.</u> The average fraction of ORF length predicted to encode disordered residue in adaptive, neutral and deleterious emerging ORFs is shown. Adaptive emerging ORFs are less disordered that neutral and deleterious emerging ORFs (Mann-Whitney U test *P*=0.03 and *P*=0.02, respectively). Error bars: s.e.m. **B.** <u>No strong association between GC content and fitness.</u> Adaptive emerging ORFs have a slightly lower GC content than neutral ones (Mann-Whitney U test *P* = 0.04). Error bars = s.e.m. <u>**C-F.** Strong association between high TM propensity and beneficial fitness effects.</u> The average fraction of ORF length predicted to encode TM residues (**C-D**) and the fraction of ORFs predicted to contain at least one full TM domain (**E-F**) according to TMHMM (**C, E**) and Phobius (**D, F**) in adaptive, neutral and deleterious emerging ORFs are shown. TM propensity is significantly greater in adaptive than neutral emerging ORFs in all cases (**C-D**: Mann-Whitney U tests *P*<0.0025; **E-F**: Fisher’s exact tests *P*<0.025). Error bars: s.e.m. (**C-D**) and standard error of the proportion (**E-F**).

The 28 adaptive emerging ORFs we identified as increasing relative fitness when overexpressed did not seem any more required for survival than other emerging ORFs (*P*>0.05 when comparing fitness cost of deletion, ORF intactness and nucleotide diversity across isolates). Furthermore, these ORFs were never found to be toxic in any of the conditions we tested, in contrast with established ORFs which can be toxic in one environment even when adaptive in another. It is thus unlikely that these adaptive emerging ORFs have already evolved useful physiological roles, in line with the adaptive proto-gene evolution prediction (Fig. 1A). Overall, our experiments show that expression of dispensable emerging ORFs can provide fitness benefits across multiple environments, and thus facilitate *de novo* gene birth.

## Beneficial biochemical capacities

Although we predicted the adaptive potential of emerging ORFs, the molecular mechanisms that may mediate such beneficial effects remain mysterious. It has been suggested that high levels of intrinsic structural disorder may be associated with adaptive fitness effects^27^ based on the observation that young *de novo* genes are highly disordered in many species^27–30^. However, in *S. cerevisiae*, recently-evolved ORFs are predicted to be less disordered than conserved ones^3, 7, 29, 31^ and increasing the expression of disordered proteins causes deleterious promiscuous interactions^32^. The 28 adaptive emerging ORFs identified in our screens did not exhibit high intrinsic disorder (Fig. 4A). In fact, their translated products were predicted to be significantly less disordered that neutral and deleterious emerging ORFs (Mann-Whitney U test *P*=0.03 and *P*=0.02, respectively). Our data thus indicates that disorder is unlikely to be a beneficial biochemical capacity that promotes *de novo* gene birth in *S. cerevisiae*, although it may be in other lineages with more complex regulatory systems^33^.

In *S. cerevisiae*, young ORFs display high GC content^7^ and TM propensity, the latter presumably mediated by a sequence composition biased towards hydrophobic and aromatic residues^3, 8, 34^. We investigated whether these properties may promote beneficial fitness effects in *S. cerevisiae*, after verifying that adaptive, neutral and deleterious emerging ORFs presented indistinguishable ORF length distribution (Mann-Whitney U tests *P*>0.3 for all comparisons). GC content was slightly lower in adaptive than neutral emerging ORFs (Fig. 4B). Adaptive emerging ORFs however displayed strikingly high TM propensity as measured by both average TM residue content and fraction of ORFs with full TM domains according to two prediction algorithms (Figs. 4C-F). Though remarkably pronounced and robust in emerging ORFs, the association between TM propensity and fitness benefits was absent in established ORFs (Extended Data Figs. 5–6). Thus, our data suggests that emerging ORFs containing TM domains promote fitness in budding yeast.

## Cryptic origins of transmembrane domains

These putative adaptive TM domains may have evolved progressively throughout the ORF emergence process (ORF-first) or may result from intergenic sequence biases present in the genome prior to ORFs emergence (TM-first). To distinguish between these scenarios, we compared the TM propensities of established and emerging ORFs with those of artificial ORFs corresponding to the hypothetical translation products of intergenic sequences predicted after removing intervening stop codons (iORFs; **Methods**). TM propensities were lowest in established ORFs, intermediate in emerging ORFs and highest in iORFs (Fig. 5A). This result, consistent with a previous study^34^, held when established ORFs and iORFs were sampled to match the length distribution of emerging ORFs (Extended Data Fig 7). Scrambled control sequences, retaining the same length and nucleotide composition as the real genomic sequences, displayed surprisingly high TM propensities (Figs. 5A). Altogether, these observations supported the TM-first scenario.

**Figure 5.**
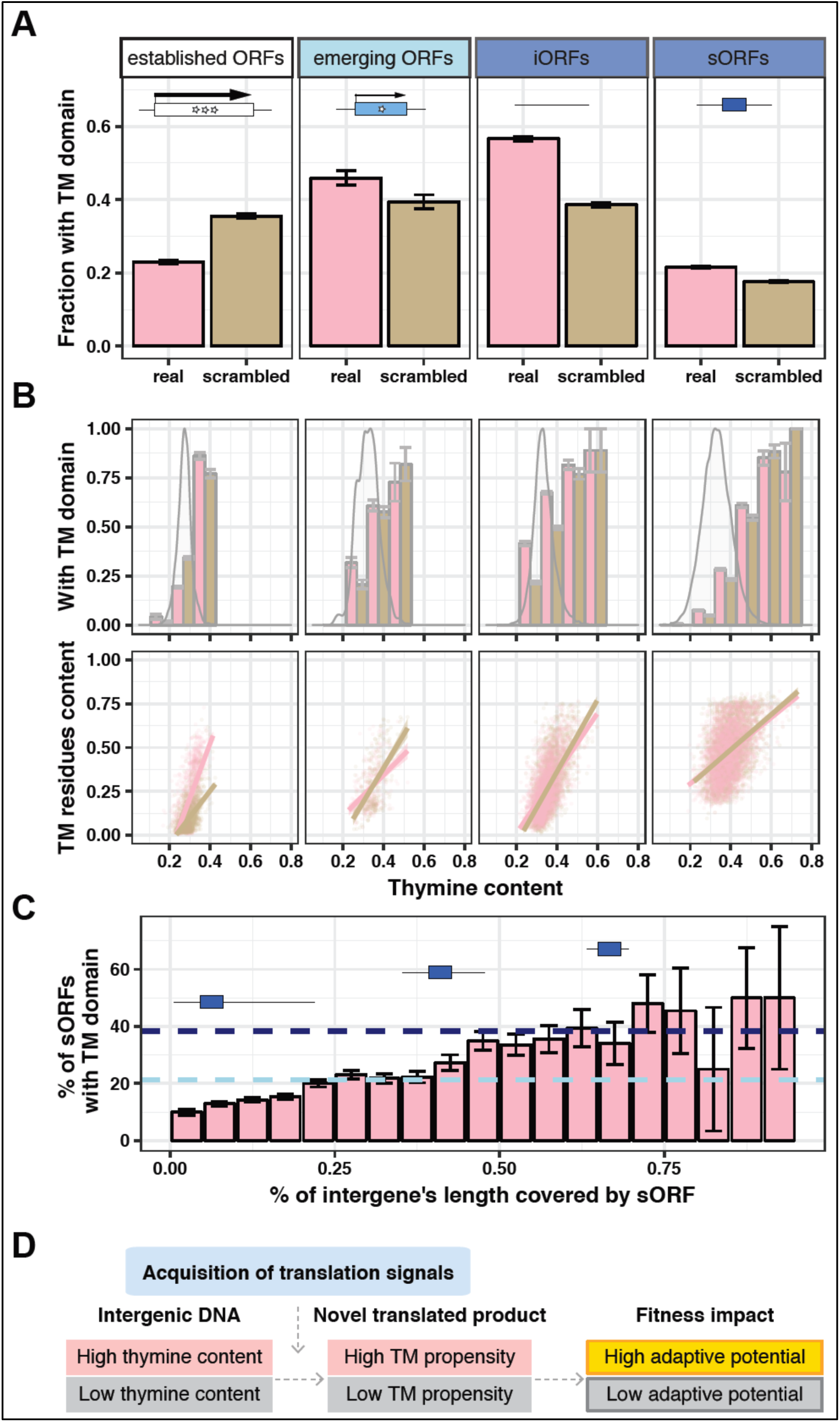
Pervasive transmembrane propensity throughout the genome. **A.** <u>Propensity of ORFs to form TM domains.</u> Scrambled sequences maintain the same length and nucleotide composition as the real genomic sequences. Intervening stop codons are removed from iORFs and scrambled sequences. sORFs occur pervasively in the genome. Error bars: standard error of the proportion. **B.** <u>Thymine content influences TM propensity.</u> From left to right: established ORFs, emerging ORFs, iORFs, sORFs. Top panel: Bar graph represents the fraction of ORFs that encode a putative TM domain, binned by thymine content (bin size = 0.1) and compared between real (pink) and scrambled (brown) sequences. Error bars represent standard error of the proportion. Overlaid density plots represent the distribution of all sequences per category. Bottom panel: scatterplot showing the fraction of sequence length predicted to be TM residues as a function of thymine content. Only ORFs predicted to encode a putative TM domain are included in the bottom panel. Individual real (pink) and scrambled (brown) sequences are shown in the scatterplot with transparency; points of higher intensity indicate that sequences sampled multiple times. Linear fits with 95% confidence intervals are shown. **C.** <u>Additional intergenic sequence signals increase the probability that sORFs contain TM domains</u>. Only sORFs between 25 and 75 codons that are fully contained within intergenes were included in this analysis. Blue rectangles on horizontal black lines illustrate how sORFs of a given length can occupy a small (left) or large (right) fraction of intergenic sequence length. Horizontal dashed lines represent the fraction of emerging ORFs (light blue) and iORFs (dark blue) between 25 and 75 codons predicted to contain putative TM domains. Error bars represent standard error of the proportion. **D.** <u>A new model for adaptive proto-gene evolution.</u> Our data suggests that intergenic thymine content influences the TM propensity of novel translated products, which in turn influences their adaptive potential.

Previous analyses encompassing multiple species have shown that GC content negatively correlates with expected TM propensity^29^ and that the TM domains of established membrane proteins consist of stretches of hydrophobic and aromatic residues encoded by thymine-rich codons^35^. We thus hypothesized that the high thymine content of yeast intergenic sequences (Extended Data Fig. 6) may facilitate the emergence of novel polypeptides containing TM domains with adaptive potential. In agreement with this hypothesis, a strong influence of thymine content on TM propensity was observed regardless of ORF type, ORF length, or whether the sequences were real or scrambled (Fig. 5B, Extended Data Fig. 7). Established ORFs displayed non-random TM propensities, while emerging ORFs and iORFs appeared enriched in TM domains relative to their thymine content (Figs. 5A-B, Extended Data Fig. 7). We also estimated the TM propensity of small unannotated ORFs that pervasively occur throughout the genome (sORFs). TM propensity in sORFs was also largely driven by their thymine content, and markedly increased when they occupied a larger portion of the intergenic region from which they were extracted (Figs. 5B-C). Altogether, these results showed that the yeast genome harbors a pervasive TM propensity, facilitated by a high thymine content, and further magnified by additional intergenic sequence signals which possibly reflect a form of preadaptation to the birth of novel proteins^36^ or other constraints. This discovery converges with our finding that overexpressing emerging ORFs with TM domains tends to increase relative fitness (Fig. 4). Together, these two findings suggest that proto-genes with adaptive biochemical capacities, such as TM domains, are most likely to emerge when the blueprint for these capacities existed in the genome before the acquisition of translation capacities (Fig. 5D).

**Fig. 6.**
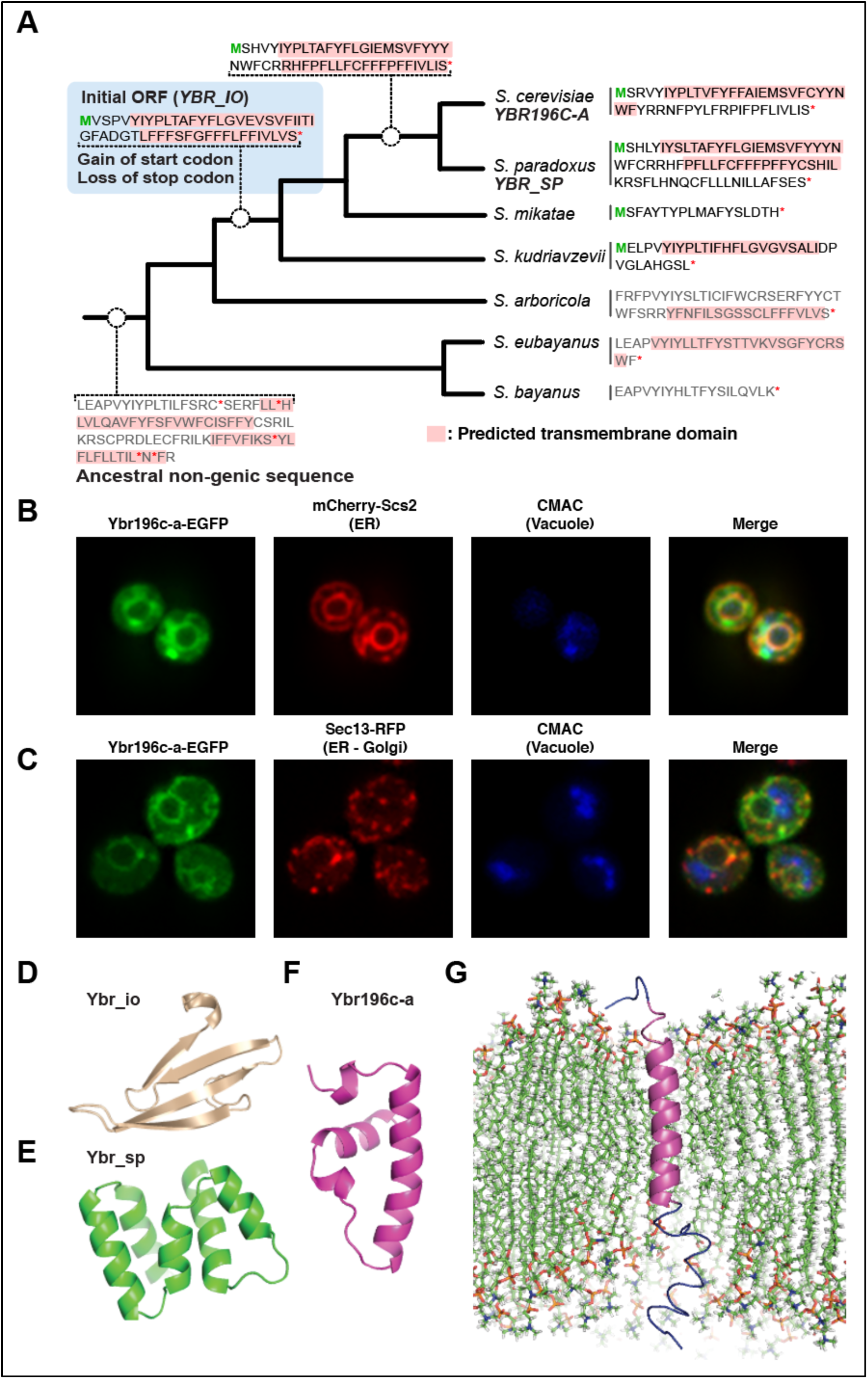
*YBR196C-A*, a TM emerging ORF with adaptive potential. **A.** <u>*YBR196C-A* emerged in an ancestral non-genic sequence with high cryptic TM propensity.</u> Results of ancestral reconstruction software are summarized along the *Saccharomyces* genus phylogenetic tree. Branch lengths are not meaningful. Theoretical translations of extant and reconstructed sequences are shown, with ORF boundaries indicated by a green M and a red *. ORF-less ancestors displayed were chosen for illustration purposes. Except for the intergenic ancestor, the translation of the sequence that aligns to the start codon of *YBR196C-A* is shown, in the relevant frame, and until the first stop codon is reached. **B.** <u>Ybr196c-a colocalizes at the ER with Scs2p.</u> Chromosomally integrated Scs2-mCherry and plasmid-borne Ybr196c-a-EGFP localization was assessed using confocal microscopy. **C.** <u>Ybr196c-a colocalizes at the ER, but not at the Golgi, with Sec13p.</u> Chromosomally integrated Sec13-RFP and plasmid-borne Ybr196c-a-EGFP localization was assessed using confocal microscopy. **D-F.** <u>Predicted 3D structures for the translation products of *YBR_IO* (**D**), *YBR_SP* (**E**), *YBR196C-A* (**F**).</u> Model 1 predictions by Robetta are shown. *YBR_IO* is predicted to encode an all-*β* strand protein. *YBR_SP* is predicted to encode a protein with multiple short *α*-helices (longest are 13 and 14 residues; between positions 18 and 31, and 58 to 72, respectively). *YBR196C-A* is predicted to encode a protein with a long *α*-helix (20 residues; positions 9 to 29 between Pro and Arg). **G.** <u>Ybr196c-a is predicted to stably integrate membranes</u>. Molecular dynamics simulation shows that, after 200ns, the peptide has kept the helix intact, with N and C terminal tails interacting with the surface of the lipid bilayer.

## An emerging ORF with adaptive potential caught in the process of fixation

To further investigate this hypothesis, we sought to retrace the evolutionary history of a specific locus to determine whether an ORF-first or TM-first scenario was most likely. We focused on *YBR196C-A*, one of the 28 adaptive emerging ORFs identified on our screens that was predicted to contain a TM domain and whose ORF structure appears relatively stabilized within *S. cerevisiae* (intact ORF in 95% isolates; one of six annotated ORFs in the genome with these characteristics). Extensive sequence similarity searches across a broad phylogenetic range (**Methods**) failed to identify sequences similar to *YBR196C-A* in species beyond the *Saccharomyces* genus, consistent with a recent origin. Aligning syntenic sequences across six *Saccharomyces* species revealed that ORFs of varying lengths in different reading frames were present in some, but not all, species of the clade, with highly variable primary sequences (Extended Data Fig. 8). Ancestral reconstruction along the clade (**Methods**; Fig. 6C) showed that no potential ORF longer than 30 codons was present in the *Saccharomyces* ancestor, in any reading frame (Extended Data Fig. 8), confirming the *de novo* origination of *YBR196C-A*.

The initial ORF that became *YBR196C-A* (*YBR_IO*) likely originated at the common ancestor of *S. kudriavzevii*, *S. mikatae*, *S. paradoxus* and *S. cerevisiae* and already encoded putative TM domains. In fact, the ancestral intergenic sequence at the base of the clade already contained a suite of codons that would have had the capacity to encode TM domains, had it not been interrupted by stop codons (Fig. 6C). This TM propensity persisted in most extant sequences despite substantial primary sequence changes. Consistent with our previous analyses (Fig. 5), *YBR196C-A* is extremely T-rich (48%, 99^th^ percentile of all annotated ORFs) and so are its extant relatives and reconstructed ancestors. The inferred evolutionary history of the *YBR196C-A* locus was therefore largely consistent with a TM-first scenario.

Besides its predicted TM propensity, there is no published evidence that the protein encoded by *YBR196C-A* has the potential to integrate into cellular membranes. The predicted TM propensity could be an artifact arising from having applied existing TM prediction methods, which were trained on established membrane proteins. To test this, we visualized cells overexpressing *YBR196C-A* tagged with green fluorescent protein by confocal microscopy (**Methods**). The protein colocalized with two markers of the endoplasmic reticulum membrane: Scs2p and Sec13p (Figs. 6A-B, Extended Data Fig. 9). In a fraction of the cells, the protein also localized to puncta, where colocalization was observed with Scs2p, but not Sec13p (Extended Data Fig. 9). We did not observe localization at the cell periphery, nor colocalization with mitochondrial, peroxisomal or vacuolar markers (Extended Data Fig. 9). The protein encoded by another emerging ORF, *YAR035C-A*, was observed specifically in the mitochondrial membrane when we visualized it using the same methods (Extended Data Fig. 10). Therefore, the *YBR196C-A* locus has the potential to encode a protein that associates with a select subset of cellular membranes.

*YBR_IO* displayed a strong TM propensity, but underwent major changes in primary sequence including frameshifts, truncations and elongation throughout *Saccharomyces* evolution (Fig. 6C). Furthermore, examination of syntenic loci in five *S. paradoxus* isolates showed that the *YBR196C-A* homologue in this species (*YBR_SP)* failed to display an equivalent level of intraspecific constraints as *YBR196C-A* in *S. cerevisiae*. An ORF was present in the syntenic loci of four out of five isolates but it displayed frameshift-induced variation in length and sequence, despite substantial conservation in the TM-containing N-terminal region of the alignment (Extended Data Fig. 8). Thus, our screening possibly “caught” *YBR196C-A* in the process of actively establishing itself in the *S. cerevisiae* genome, after going through substantial changes since *YBR_IO* first presented blueprints for TM domains to the action of natural selection.

We further determined that adaptive mutations are actively shaping the molecular changes observed in the protein sequence, consistent with our model (Fig. 1). A positive selection test across the four *Saccharomyces* species containing ORFs yielded statistically significant results (*P = 0.01*, LRT: *D* = 8.9, *df* = 2) and identified three sites under positive selection. These results were robust to the choice of statistical model (**Methods**) and showed slightly increased significance when focusing on the TM region of the alignment (First 30 codons: P = 0.007, LRT: D = 10, df = 2). Furthermore, a pairwise d_N_/d_S_ (omega) calculated over the first 30 codons of the alignment between *S. cerevisiae* and *S. paradoxus* was significantly greater than 1 regardless of the model used (YN: 4.5, LWL85: 1.7, LPB93: 3.8, NG86: 1.9).

To investigate the impact of this rapid sequence evolution at the protein level, we compared the predicted 3D structures of the putative proteins encoded by *YBR_IO*, *YBR_SP* and *YBR196C-A*. All three shared a conserved predicted TM domain in the N-terminal region, but this domain contained a Gly in *YBR_IO* and *YBR_SP* that mutated to Ala in *YBR196C-A* (Extended Data Fig. 8). A second, Phe-rich, low complexity TM domain was predicted for *YBR_IO* and *YBR_SP* in the C-terminal region of the alignment, again presenting Gly and Pro residues (Extended Data Fig. 8). Gly and Pro are known for their low helix propensity in solution^37^ and in a membrane environment^38^, and expectedly hindered the formation of *α*-helical structures in 3D models generated with the *ab initio* prediction server Robetta^39^. In fact, the models were almost all *β* strands rather than helices for *YBR_IO* (Fig. 6D). Models of *YBR_SP* displayed short, broken *α*-helices relative to models of *YBR196C-A*, due to the extra Gly and Pro residues (Figs. 6E, F; Extended Data Fig. 8). The only structural models capable of spanning the membrane were those for *YBR196C-A*. Four out five models predicted by Robetta for Ybr196c-a yielded a robust TM helical structure spanning 20 residues flanked by Pro in N-terminal and a pair of charged Arg in C-terminal, and explicit solvent molecular dynamics simulations of these 3D structures in a membrane bilayer strongly supported that *YBR196C-A* has the potential to encode a single pass membrane spanning protein (Fig. 6G).

## Discussion

Several theories predicted that emerging sequences must carry adaptive potential for *de novo* gene birth to be possible^2, 3, 40^. Our results validate this prediction and support an experiential model for *de novo* gene birth whereby a fraction of incipient proto-genes with adaptive potential can subsequently mature and, as adaptive changes engender novel selected effects, progressively establish themselves in genomes in a species-specific manner (Figs. 1, 6). This model, consistent with studies showing that *de novo* emerging sequences remain volatile for millions of years^5, 22, 41–44^, emphasizes that the selective pressures acting on proto-genes are different from those acting on canonical protein-coding genes.

Our work suggests that incipient proto-genes that expose cryptic TM domains through translation are more likely to carry adaptive potential than others, and thus more likely to contribute to evolutionary innovation through positive selection in *S. cerevisiae* (Figs. 4–6). This novel mechanistic model may explain how sequences that were never translated previously can evolve into novel proteins with useful physiological capacities. Indeed, we show that a simple thymine bias in intergenic sequences suffices to generate a diverse reservoir of novel TM peptides (Fig. 5). The membrane environment might provide a natural niche for these novel peptides, shielding them from degradation by the proteasome, and allowing subsequent evolution of specific local interactions while reducing the potential for deleterious promiscuous interactions throughout the cytoplasm. Our examination of the *YBR196C-A* locus illustrates how a thymine-rich intergenic sequence with high TM propensity can, upon acquisition of translation signals, be molded by positive selection into a genuine TM ORF with adaptive potential that acquires selected effects as it matures over millions of years.

Our discovery that emerging ORFs containing TM domains tend to increase fitness when overexpressed (Fig. 4) is surprising given the elaborate systems controlling the insertion and folding of TM domains and preventing their aggregation^45^. No current model could have anticipated this and, to our knowledge, this is the first report of a biochemical protein feature being empirically associated with adaptive potential besides life-saving antifreeze glycoproteins in polar fishes^46^. How might expression of a TM proto-gene confer a growth advantage? This is an exciting unanswered question raised by our studies, for which one might consider several speculative models. Expression of TM proto-genes may cause a preconditioning stress, modestly inducing the unfolded protein response (UPR), heat-shock protein or other protein chaperone expression. This type of preconditioning stress has been shown across species to confer a benefit to subsequent exposure to stressors^47–51^. Novel TM domains might also beneficially associate with larger membrane proteins, such as transporters or signaling proteins^52^. Alternatively, insertion of proto-gene TM domains could alter the biophysical properties of the lipid bilayer itself, slowing diffusion of lipids within the bilayer, reducing diffusion of molecules across the membrane or altering membrane curvature through molecular crowding^53–55^. It is important to note that no single model may explain their adaptive effects universally. How the emerging TM proteins are addressed at the membrane by the cellular machinery, or why TM domains would become selected against with evolutionary time, is as of yet unclear.

A previous analysis of young *de novo* genes in 17 *Saccharomycotina* genomes found them to be consistently more TM rich than conserved genes^34^. Furthermore, among the rare *de novo* genes with characterized phenotypes, several are predicted to encode TM domains. These include *Saturn*, a *de novo* gene specific to the fruit fly *Drosophila melanogaster* essential for spermatogenesis and sperm function^58^, and *TMRA*, a vertebrate *de novo* gene specific to the songbird Zebra finch expressed in the part of the brain associated with vocal learning^59^. The human Papillomavirus type 16 E5*α* onco-gene, which encodes a TM protein localized at the Golgi apparatus, might also have emerged *de novo*^56, 57^. The positive growth impact of *de novo* emerging TM sequences we observed in yeast might have oncogenic implications, since experiments showed that random TM sequences have the ability to transform mouse cells^17^. An abundance of cryptic TM domains in unannotated translated sequences was reported in *Drosophila*^60^ and in *E. coli*, where many novel TM proteins appear involved in stress response^61, 62^. Mammalian micro peptides translated from lncRNAs and active in muscles also have TM domains, which can act as co-factors in complex molecular processes^52, 63, 64^. Thus, the implications of cryptic TM domains in *de novo* gene birth likely generalize beyond *S. cerevisiae*.

## Methods

### Yeast strains used for screening

Barcoded haploid yeast overexpression strains in the BY4741 background from the BarFLEX collection^24^ were used for screening purposes together with the reference strain with matched genetic background. The reference strain was created by growing a randomly selected BarFLEX strain on SC+GLU+G418+5FOA to chase the plasmid, then transforming the plasmid-less strains with the destination plasmid pBY011.

Overexpressed ORFs included in the screen analyses are listed in **Source Data 1**. All strains were kept in SC-URA+GLU+G418 glycerol stocks at −80°C in 384-well format until used for screening. Growth medium composition are detailed in Extended Data Table 1, strain genotypes in Extended Data Table 2 and plasmids in Extended Data Table 3.

### Yeast strains used for microscopy

Using the Gateway System® (Life Technologies, Carlsbad, CA), the emerging ORFs (*YBR196C-A* and *YAR035C-A*) were first cloned into donor vector pDNOR223 using BP recombination (Gateway BP Clonase II Enzyme Mix, Life Technologies) and then transferred to destination vector pAG426GAL-ccdB-EGFP by LR recombination (Gateway LR Clonase II Enzyme Mix, Life Technologies). The expression vectors were then used to transform multiple strains carrying organelle markers that were chromosomally-tagged with fluorescent proteins, using the LiAc/PEG/ssDNA protocol^65^ and selected on media lacking uracil. See Extended Data Tables 2 and 3 for detailed information about the plasmids and strains used.

### Classification of S. cerevisiae ORFs

Annotated ORFs were assigned to the established group or the emerging group based on their interspecific conservation levels (young, old) and protein-coding signatures. Genome annotations for 6,310 *S. cerevisiae* ORFs from the Saccharomyces Genome Database (SGD)^66^ were used keeping a consistent version with ref^3^. Protein-coding signatures were derived from 3 lines of evidence as in ref^3^: signatures of translation in ribosome profiling data sets; signatures of intra-species purifying selection estimated from 8 *S. cerevisiae* strains; ORF length longer than expected by random mutations (300nt). ORFs were categorized as young when they 1) had no homologues detectable by phylostratigraphy beyond the *Saccharomyces* clade (conservation level *≤*4) as in^3^ and 2) had no syntenic homologue with more than 50% sequence similarity in *S. kudriavzevii* or *S. bayanus* (see *Synteny analysis* below). All young ORFs that displayed less than 3 of the protein-coding signatures were assigned to the emerging ORFs group. All other ORFs (old ORFs and young ORFs with strong protein-coding signatures) were assigned to the established ORFs group.

We also defined artificial intergenic ORFs (iORFs) and genomic small ORFs (sORFs) as follows. iORFs were generated as in ref^7^: first, non-annotated genomic regions were extracted using *bedtools subtract* ^67^ and the annotation GFF file downloaded from SGD. Then, stop codons in the +1 reading frame were removed, and the sequences in that frame were translated and used to calculate the various properties. sORFs include all non-annotated ORFs that showed no signs of translation from ref^3^, that were longer than 75nt, and did not entirely overlap annotated ORFs in any strand (n=18,503).

### Synteny analysis

We identified syntenic blocks for young ORFs across four *Saccharomyces* species by using the downstream and upstream genes of the young ORFs as anchors The orthologs of the anchor genes were downloaded from ref^68^. In cases where a continuous syntenic block could not be constructed between two anchor genes, it was constructed by aligning the ±1kb region surrounding the anchor gene with the largest number of orthologs (identified by ref^69^; downloaded from SGD). Multiple alignment of the syntenic blocks were generated using MUSCLE^70^ using the default parameters of the msa R package^71^.

For each syntenic block, we identified all non-*S. cerevisiae* ORFs that overlapped with the *S. cerevisiae* young ORFs within the ORF±200bp region of the alignment. The overlapping region of the ORFs were extracted from the alignment, translated, and re-aligned pairwise. Similarity between ORFs was calculated by taking the ratio of the number of identical amino acids over the overlapping portion of the alignment divided by the length of *S. cerevisiae* ORF. All young *S. cerevisiae* ORFs with similarity higher than 50% with ORFs in *S. bayanus*, and *S. kudriavzevii* were excluded from the list of emerging ORFs and considered established instead.

### Annotation status of emerging ORFs

At least 95% of emerging ORFs are annotated as dubious or uncharacterized according to both the 2011 *S. cerevisiae* genome annotation (consistent with^3^) and the current one (R64-2-1). Both annotations were downloaded from SGD.

### Primary sequence analyses

Estimates for intrinsic disorder were lifted from^3^. GC content was calculated using a Python script. Percentage of residues in transmembrane domains and number of transmembrane domains were predicted using the online servers of TMHMM^72^ and Phobius^73^ (**Source Data 1**) and secondary structure for extant and reconstructed *YBR196C-A* homologues was predicted using psiPred^74^ (Extended Data Fig. 8).

### Genome-wide overexpression screening on ultra-high-density colony arrays

Using a Singer ROTOR robotic plate handler (Singer Instrument Co. Ltd), overexpression and reference strains where transferred from glycerol archives to agar plates and then robotically combined into 1536-density agar plates (SC-URA+GLU+G418; in Extended Data Table 1). Cells on these plates were then transferred with the same robot at the same density to SC-URA+GAL+G418 where they were incubated for a day. This process was repeated once, following which the cells were robotically transferred to 6144-density agar plates in the screening conditions (in Extended Data Table 1). Throughout this transfer process, five 1536-density source plates were copied 4 times, yielding on average 4 technical replicates per overexpression strain and 3,072 replicates for the reference strain per screening condition. Specifically, 768 replicates of the reference strain were arrayed on one out of the five 1536-density plates in an alternating pattern with a replicate of the reference strain in every other row/column. Only four 1536-density source plates can be transferred to a single 6144-density plate, therefore 4 out of the five 6144-density plates contained 768 replicates of the reference strain. The alternating pattern of the reference strain from the 1536-density plate was carried over to the 6144-density plate. As a result, the reference strain was systematically spread throughout the plate, appearing every 4 columns/rows, and each colony replicate of the same overexpression strain was always surrounded by different neighbors. This experimental set up was designed to mitigate neighbor effects, for instance a fast-growing colony negatively impacting the growth of its neighbors.

### Quantification of colony sizes

Colony size was estimated at the time point when the reference strain had reached saturation, which varied depending on the screening condition (**Source Data 3**). Digital images of the plates were acquired in 8-bit JPEG with a SLR camera (18Mpixel Rebel T3i, Canon USA Inc., Melville/NY) with an 18-55 mm zoom lens. A white diffusor box with bilateral illumination and an overhead mount for the camera in a dark room were used. The workflow we used for colony size quantification was similar to^75^: (a) each plate was imaged 3 times to control for technical variability in image acquisition, (b) each image was cropped to the plate, (c) a colony grid at expected density was overlaid on the plate image, (d) each image was binarized by thresholding the ratio between background pixel intensity vs. colony pixel intensity, (e) the number of pixels at each position on the grid was counted, (f) empty positions on the grid were temporarily removed to avoid affecting the spatial normalization in the next steps, (g) the number of pixels at each position on the grid were averaged across all 3 images, (h) spatial normalization and border normalization were applied to remove local, nutrient-based growth effects as in^76^, (i) grid values were normalized by the mode of pixel counts per plate to allow comparison of values across plates, (j) empty positions on the grid were added back, (k) outer rows/columns were excluded from downstream analysis (depth of 5 for 6144-density plates) because, even after applying the spatial and border normalization algorithms, reduced border artifacts could still be detected. Normalized colony size was then defined as the average of the normalized number of pixels per grid point corresponding to technical replicates of each overexpression strain.

### Identification of overexpression strains with increased and decreased relative fitness

For every screen, we binned all screened ORFs in 3 categories (increased fitness, decreased fitness and unchanged) according to their relative fitness compared to the reference strain. The *P*-values of a non-parametric Mann-Whitney U test comparing the distributions of technical replicates of each overexpression strain with that of the reference strain were corrected for multiple hypothesis testing using *Q*-value estimations as defined in^77^. ORFs where categorized as having increased or decreased relative fitness effect when overexpressed when the *Q*-value was lower than 0.01 and the normalized colony size was higher than 95%, or lower than 5%, of the technical replicates reference, respectively. All other ORFs were classified as unchanged.

The results of the five screens presented in Fig. 3 are reported in **Source Data 3**. These results were integrated as follows: ORFs that increased fitness when overexpressed in at least 1/5 conditions were labelled “adaptive”; ORFs that decreased fitness in at least 1/5 conditions and never increased it in any of the 5 conditions were labelled deleterious; all other tested ORFs were labelled neutral.

### Simulated ORFs for transmembrane propensity analyses

Scrambling of nucleotide sequences was performed with a custom Python script, as follows: nucleotide positions of the sequence were randomized and whenever an in-frame stop codon was formed from the randomization, its 3 positions were randomized again until they did not form a stop codon. Sequences analyzed in Fig. 5 and Extended Data Figs. 6–7 were at least 25 codons long.

In order to obtain a length distribution from a target population (e.g. established ORFs) similar to a template population (emerging ORFs), we performed sampling with replacement using a version of inverse transform sampling as follows. First, the template population’s distribution (the emerging ORF length distribution in our case) was calculated using bins of 25 codons, grouping those few of length between 300 and 650 codons. One thousand ORFs from the target populations were drawn with replacement according to this distribution. For sORFs specifically, all instances between 150 and 650 codons were grouped together as long sORFs are extremely rare. The data used to perform these analyses are shared in **Data S4** and **Data S5**.

### 3D structure predictions for extant and reconstructed *YBR196C-A* homologues

Since these sequences showed no significant homology with known proteins, an *ab initio* prediction server, Robetta^39^, was used for 3D structure predictions. Translated sequences for *YBR196C-A*, *YBR-SP* and *YBR_IO* were used as an input for *ab initio* prediction. For each sequence, 4 out of 5 model structures showed high structural similarity with slight differences in the conformation of the N and C-terminal (Model 1 for each sequence is shown in Figs. 6D-F). Ybr196c-a with the predicted TM domain spanning the membrane and the C-terminal beyond R30R31 outside the membrane was further simulated using molecular dynamics (Fig. 6G).

### Molecular dynamics simulation protocol

Membrane simulation inputs were generated using CHARMM-GUI^78, 79^. Molecular dynamics were run using CHARMM36m^80^ force field parameters at 303.23 K using Langevin dynamics. The cell box size varied semi-isotropically with a constant pressure of 1 bar using Monte Carlo barostat. Six-step CHARMM-GUI protocol for 225ps was used for equilibration. Particle Mesh Ewald (PME) was used for periodic boundary conditions (PBC) for evaluation of long-range electrostatic interactions, Lennard-Jones force-switching function used for van der Waals (VDW) and electrostatics calculations with nonbonded cutoff 12 Å. Simulations were run with 2 fs time-step utilizing SHAKE algorithm to constraint hydrogen bonds. All simulations were run for 200 ns using Ambertools 18^81^ with Cuda and first 50ns were disregarded as equilibration time. DPPC^82^ was chosen for building the lipid bilayer as PC is highly abundant in yeast membranes^83^ and bilayer thickness of around 37 Å is consistent with that of the ER membrane^84^. TIP3P explicit water model, KCl with 0.15 M and default water thickness of 17.5 Å were used. Length of X and Y was taken as 75 Å and DPPC ratio was used 10:10, resulting in approximately 83 lipid molecules on both leaflets of the membrane. Terminal group patching applied to N and C terminals of the peptide. The initial position of the TM region of the peptide was oriented using aligned along the Z axis. The system built with replacement method and ions added with Ion Placing method of “distance”. Visual Molecular Dynamics^85^ and PyMol^86^ version 1.8 were used for analysis and visualization of the trajectories.

### Intraspecific evolutionary analyses

The VCF file for 1011 *S. cerevisiae isolates* sequenced by^21^ was downloaded from the 1002 Yeast Genome website (http://1002genomes.u-strasbg.fr/files/1011Matrix.gvcf.gz). For each single-exon annotated ORF, nucleotide diversity and ORF intactness (defined as presence of start codon and stop codon in the same reading frame, with no intermediate stops) was derived from the VCF file based on all isolates that had calls for every position in the sequence using custom scripts.

Genomes of five *S. paradoxus* isolates (CBS432, N44, UWOPS91917, UFRJ50816, YPS138) were acquired from^87^, selected to give a broad representation of *S. paradoxus* evolutionary diversity. The syntenic region of *YBR196C-A* was obtained by aligning the sequence between and including the neighboring genes *YBR196C* and *YBR197C* among all *S. paradoxus* strains and *cerevisiae* S288C using MUSCLE^70^.

### Interspecific evolutionary analyses of *YBR196C-A*

Similarity searches using the Ybr196c-a protein sequence as query were performed with TBLASTN against the *Saccharomyces* genomes and all fungal genomes downloaded from GENBANK, and with BLASTP against the NCBI nr database, using a relaxed E-value threshold of 0.001 ^88^.

To reconstruct the ancestral state of *YBR196C-A*, we first identified and extracted its orthologous regions in all other *Saccharomyces* species. We exploited SGD’s fungal alignment resource to download ORF DNA + 1 kb up/downstream for guiding analyses. A multiple alignment of these sequences was generated using MAFFT^89^. A second, codon-aware alignment was generated with MACSE^90^. Using the MAFFT alignment, a phylogenetic tree was generated with PhyML^91^ with the following parameters “-d nt -m HKY85 -v e -o lr -c 4 -a e -b 0 -f e -u species_tree.nwk” where “species_tree.nwk” is the species topology. Ancestral reconstruction was performed with PRANK^92^ (on an alignment performed by PRANK, and not the one generated by MAFFT) using the above-mentioned tree as a guide and the parameters “*-showanc -showevents – F”*. The ancestral sequences were extracted from the alignment output file of PRANK, and gaps were removed to obtain the nucleotide sequences, which were then translated into amino acid sequences.

Pairwise dN/dS (omega) was calculated using yn00 from PAML^93^. Selection tests were performed using *codeml* from PAML. Specifically, the aforementioned codon-aware alignment (regenerated with only the 4 species Skud, Smik, Scer, Spar) and the corresponding PhyML guide tree were used together with the site model to perform the M1a – M2a (model=0, nsites=1 and 2) and M7 – M8 (model=0, nsites=7 and 8) Likelihood Ratio Tests of positive selection, as detailed in the PAML manual. The Bayes Empirical Bayes method at P>0.99 was used to identify sites under selection.

### Confocal microscopy

Cellular localization was determined using yeast transformed with C-terminally, EGFP-tagged emerging ORFs (see in Extended Data Table 2) expressed from the *GAL10pr* and imaged using a Nikon (Tokyo) Eclipse Ti inverted swept-field confocal microscope (Prairie Instruments, Middleton, WI) equipped with an Apo100x (NA 1.49) objective. Pre-cultures were made in SC-URA+GLU+G418 and then transferred to SC-URA+GAL+G418 for 24h, prior to imaging. Images were acquired using an electron-multiplying charge coupled device camera (iXon3; Andor, Belfast, United Kingdom) and Nikon NIS-Elements software was used to manipulate image acquisition parameters and post-acquisition processing was done using this same software, ImageJ (National Institutes of Health) and Photoshop (Adobe Systems Inc., San Jose, CA). An unsharp mask was applied in Photoshop to all images. Co-localization of EGFP-tagged emerging ORFs with subcellular membrane-bound organelles was assessed using mCherry-Scs2-TM as an ER-localized marker (**Table S2**), Sec13-RFP as a ER-Golgi marker (**Table S2**), Pex3-RFP as a peroxisomal marker (**Table S2**), MitoTracker^TM^ Red CMXRos (Thermo Fisher Scientific, Waltham, MA) mitochondrial superoxide indicator as a mitochondrial marker, and CellTracker Blue CMAC (7-amino-4-chloromethylcoumarin) dye (Life Technologies, Carlsbad, CA) as a vacuole lumen marker. Cells were incubated with 0.1 μM MitoTracker Red CMXRos for 20 min or 100 μM of CMAC blue for 15 min for mitochondrial or vacuolar staining, respectively, prior to imaging^94^. To verify that the EGFP signal was generated by our plasmid construct, we also visualized the mCherry-Scs2-TM, Sec13-RFP and Pex3-RFP strains in SC+GAL+G418 after a pre-culture in SC+GLU+G418.

## Data and materials availability

Data is available in the main text, in the extended data figures and tables, and in source data files 1-5 on github: https://github.com/annerux/AdaptiveTMproto-genes. Strains are available upon request.

## Code availability

Image processing and relative fitness estimations: https://github.com/bbhsu/protogene-analysis.

Synteny analyses: https://github.com/oacar/synal. Other analyses: https://github.com/annerux/AdaptiveTMproto-genes; Fisher’s and Mann-Whitney tests are two-sided.

## Acknowledgments

The authors are grateful: to Dr. Brenda Andrews for sharing the BarFLEX collection of overexpression strains; to Kate Licon and Dr. Shelly Trigg for technical help in handling mutant collection storage; to Dr. Philip Jaeger for help generating the reference strain used in the screens, and to UCSD undergraduates Cameron Hines, Nicholas Regent and Manuel Michaca who helped perform screens; to Drs. Graeme Sullivan and Jeffrey Brodsky for discussions; and to Drs. Gilles Fisher, Christian Landry, Benoit Charloteaux and Zoltan Oltvai for reviewing the manuscript prior to submission. This work was supported by: funds provided by the Searle Scholars Program to A-RC; the National Institute of General Medical Sciences of the National Institutes of Health grants R00GM108865 (awarded to A-RC), P41GM103504 (awarded to TI), 5R01GM097084 (awarded to CJC) and F32GM129929 (awarded to BVO); the National Institute of Environmental Health Sciences of the National Institutes of Health grant R01ES014811 (awarded to TI); the National Science Foundation MCB CAREER grant 1902859 (awarded to AFO); funding from the European Research Council under the European Union’s Seventh Framework Programme (FP7/2007–2013)/European Research Council grant agreement 309834 (awarded to AM).

## Author contributions

Conceptualization: A-RC, TI, NV; Methodology: A-RC, BH, TI, AFO, NV, AW, OA, CJC; Investigation: A-RC, NV, BH, OA, BVO, AW, JI, NCC, KM-E, AFO; Writing-Original Draft: A-RC, NV; Writing-Review and Editing: A-RC, NV, BVO, TI, AFO, BH, AW, AM, NCC, KM-E, JI, OA, CJC; Supervision: A-RC, TI, AFO, AM, CJC.

## Author Information

Authors declare no competing interests. Correspondence and requests for materials should be addressed to

---

**Extended Data Fig. 1.**
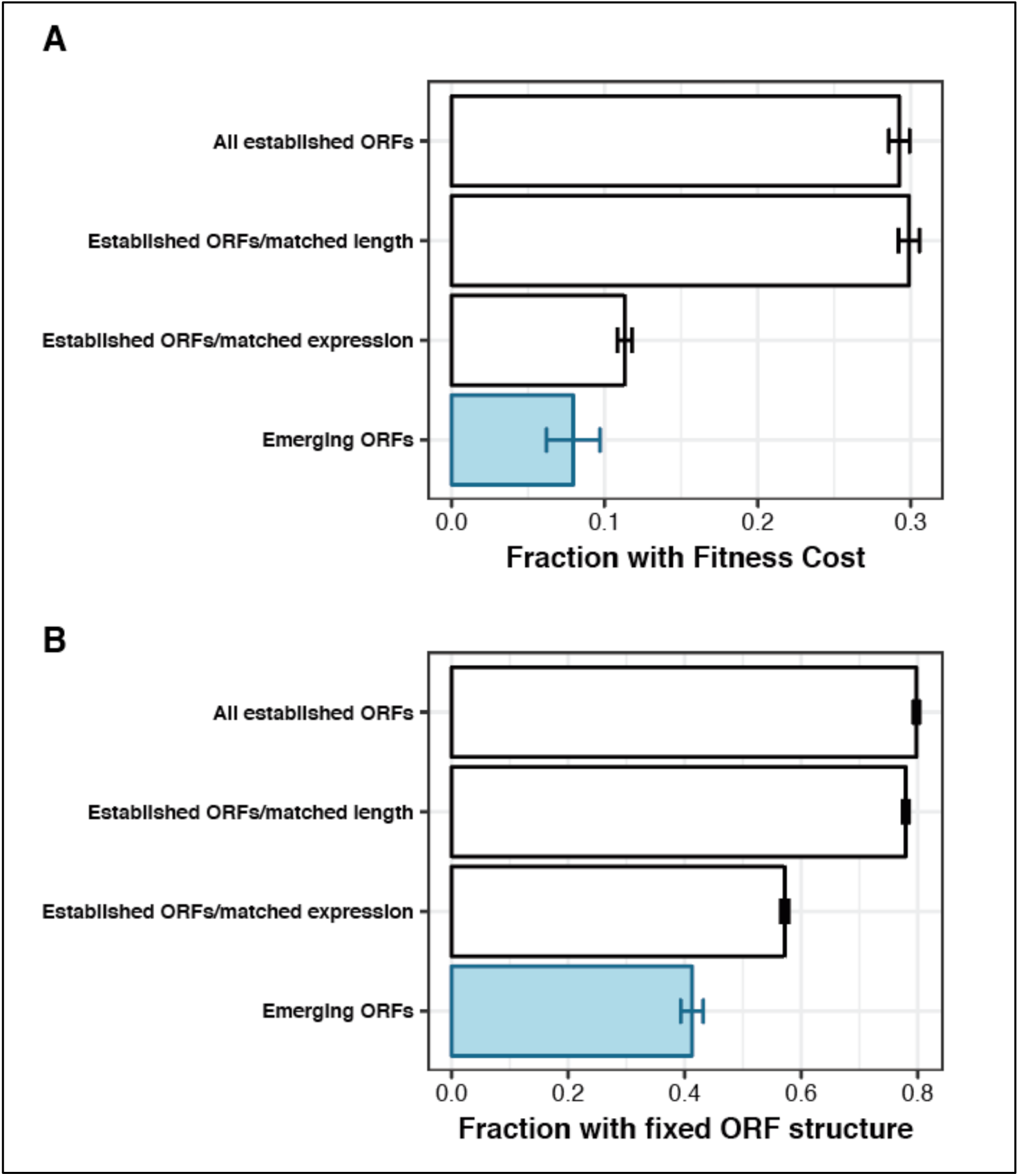
Comparing the fitness impact of loss of emerging ORFs and loss of established ORFs with matched expression levels and length distributions. Established ORFs with length and expression level distributions matched to those of emerging ORFs were randomly sampled with replacement among the set of all established ORFs (**Methods**). **A.** Fraction of ORFs for which experimental deletion leads to colonies with fitness <0.9, as in Fig. 2A **B.** Fraction of ORFs with fixed ORF structure in 90% of S. cerevisiae isolates analyzed in Fig. 2B

**Extended Data Fig. 2.**
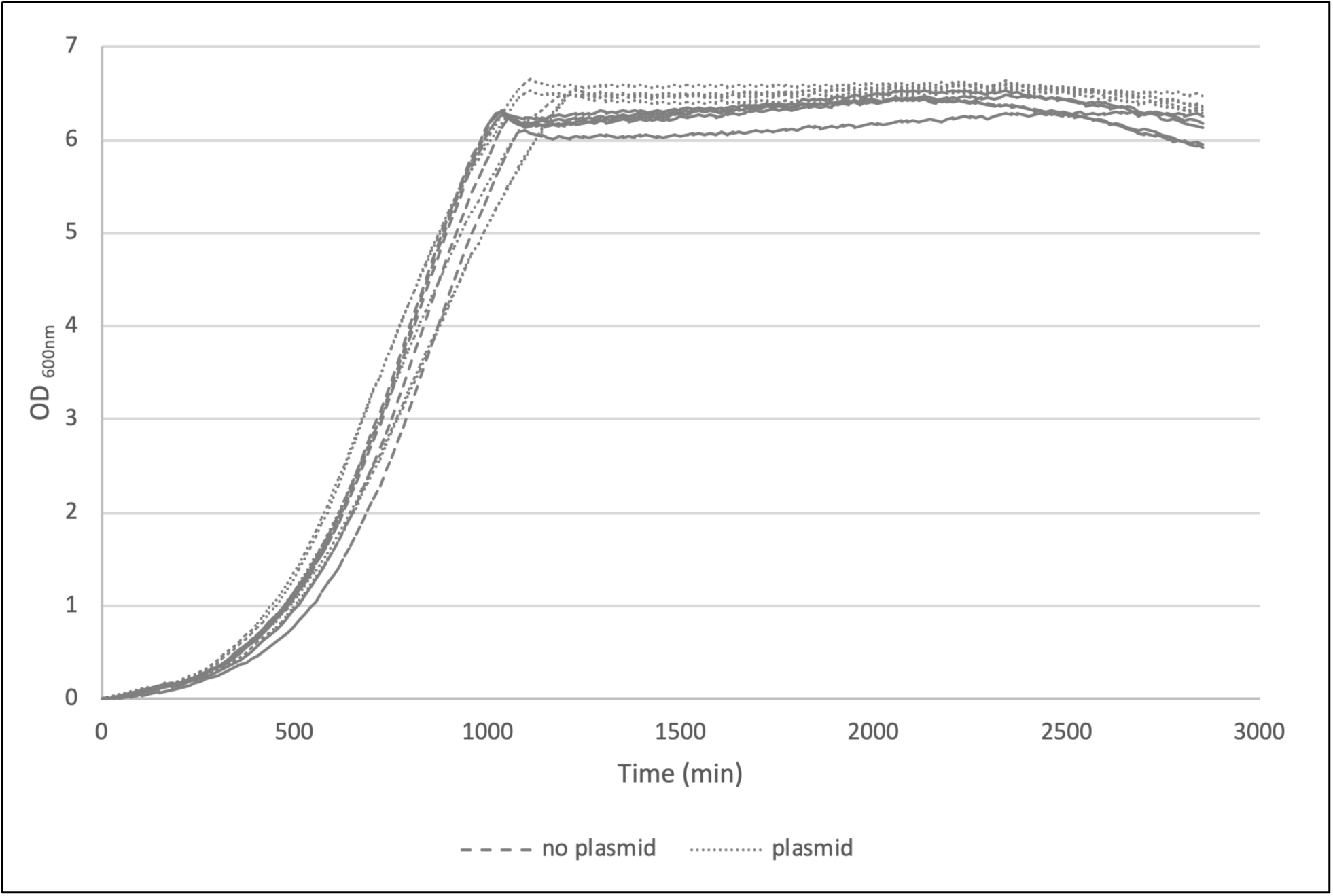
No detectable growth defect in the neutral reference strain. The reference strain was tested for growth defect by using the barcoded haploid yeast overexpression strain (**Table S2**) transformed with expression vector (pBY011) (“plasmid”) and the same strain without vector (“no plasmid”). The strains were grown in SC-URA+GAL+G418 (**Table S1**). A flat bottom 96-well plate was filled with replicates of diluted pre-cultures (125μl at OD_600nm_ 0.08) and used to collect OD_600nm_ points every 15 min for 48h using a plate reader (SpectraMax M2, Molecular Devices). For each strain, 5 biological replicates represented by 4 technical replicates each were included. Raw data was blank corrected to the respective media, path length corrected and normalized to time zero.

**Extended Data Fig. 3.**
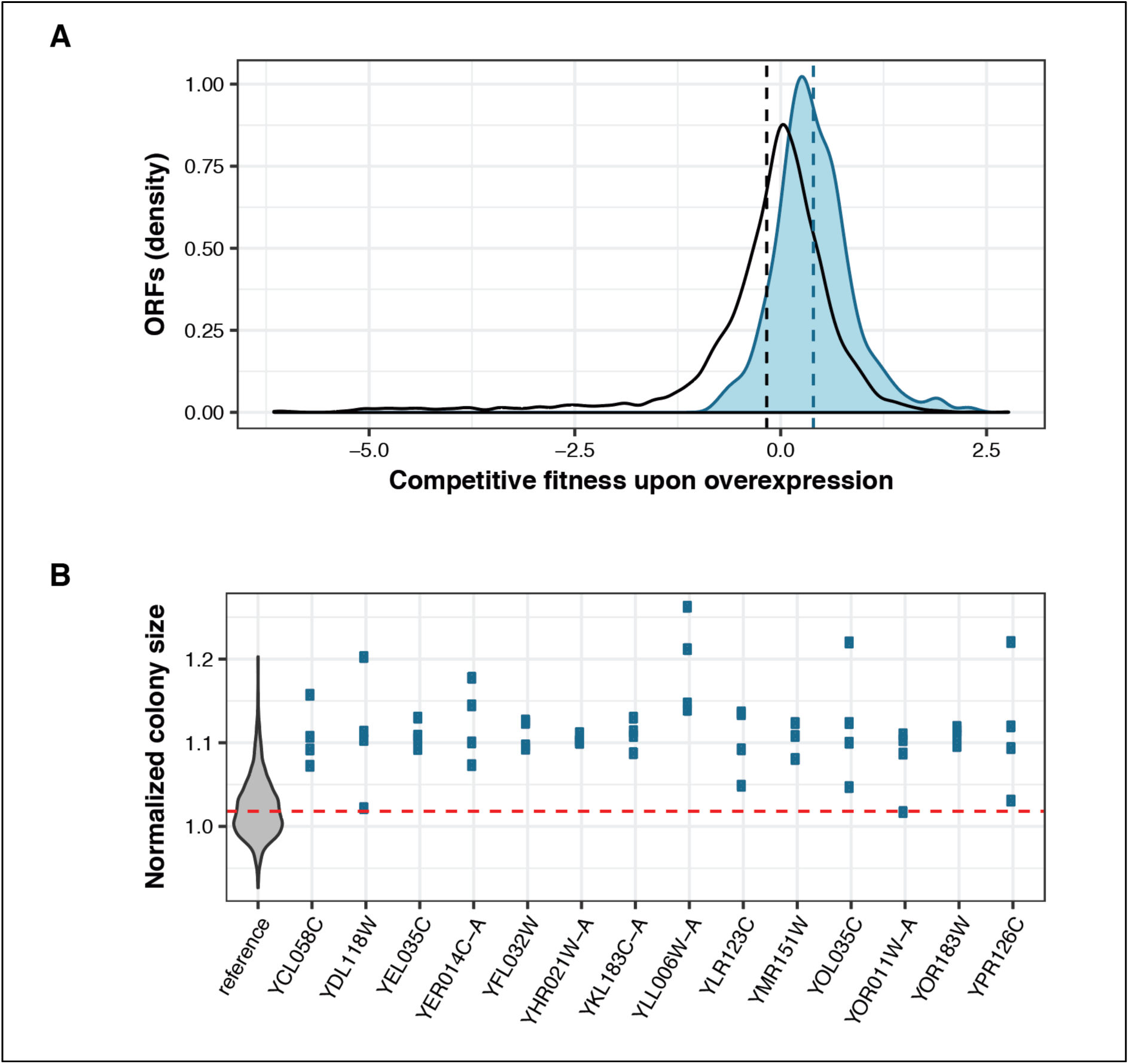
Overexpression of emerging ORFs can provide fitness benefits. **A.** <u>Emerging ORFs display higher competitive fitness when overexpressed than established ORFs (Mann-Whitney U test *P* = 5.5×10^−32^).</u> Density distributions for emerging (blue) and established (black) ORFs competitive fitness measurements in complete media after 20 generations, as measured and quantified through barcode signal intensity by^24^. Vertical dashed lines represent group means. Note that this experimental design did not allow for direct comparison with the fitness of a reference strain. **B.** <u>Distribution of the normalized sizes of individual replicate colonies for the reference strain (grey violin plot on the left) and for each emerging ORFs identified as statistically increasing fitness relative to the reference in complete media</u> (see Fig. 3B). Red dashed line represents the median normalized colony size of the reference strain. Each of the 14 emerging ORFs represented here present distributions of normalized colony sizes that are incompatible with the null hypothesis according to which they could have been randomly picked from the reference distribution (and hence were detected by our pipeline as showing increased relative fitness).

**Extended Data Fig. 4.**
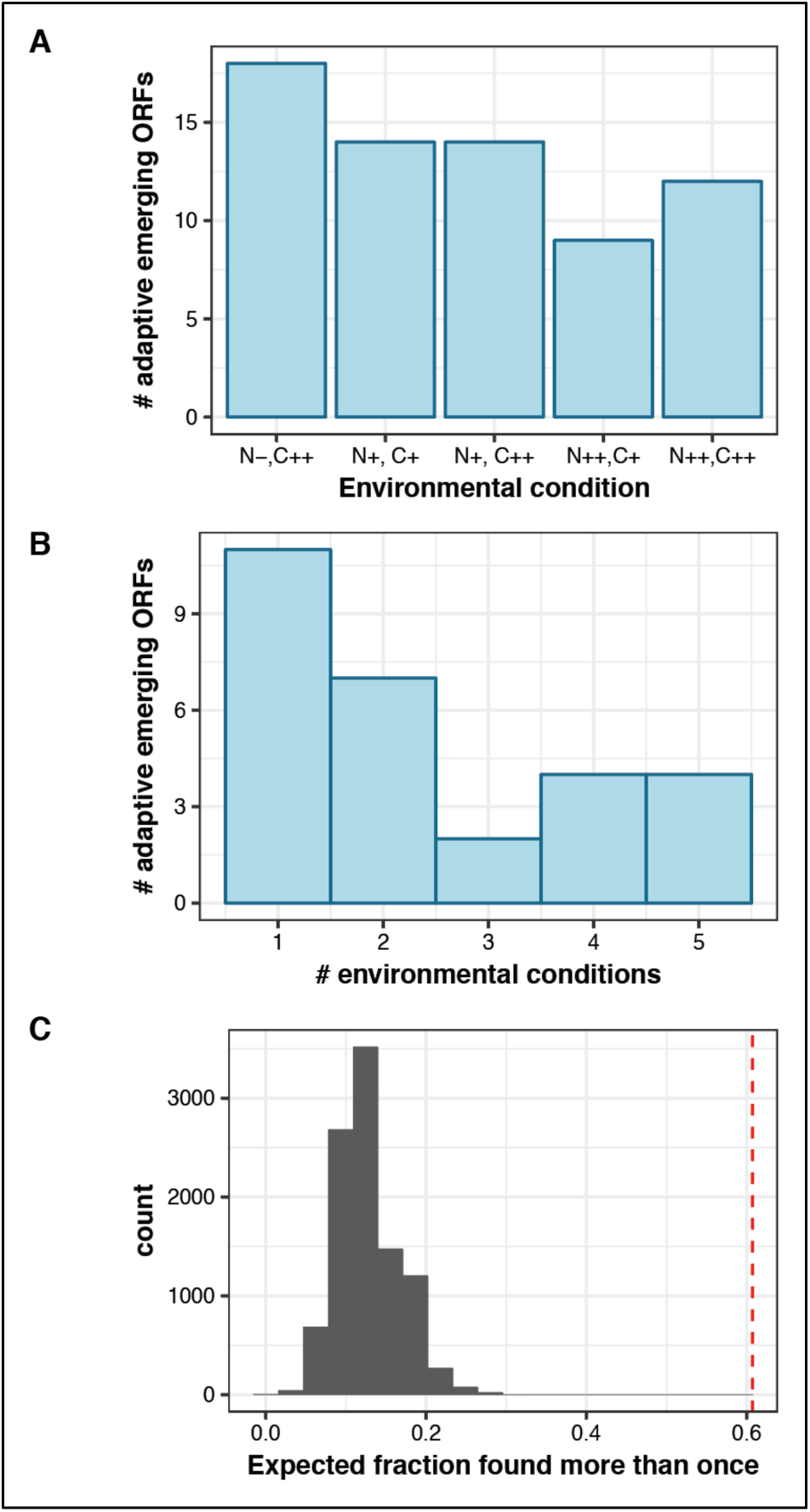
Emerging ORFs can increase relative fitness across environments. **A.** <u>Number of emerging ORFs detected as increasing relative fitness in 5 environmental conditions.</u> See **Table S1** for environment composition details. **B.** <u>Most emerging ORFs found to increase relative fitness in at least one environment are also found to increase fitness in another environment.</u> Of 28 adaptive emerging ORFs, 11 were detected in a single environment and 17 (60%) were found more than once. **C.** <u>Expected fraction of emerging ORFs that would be found to increase relative fitness in more than one environment under a stochastic null model</u> where ORFs are drawn randomly from the set of emerging ORFs never found deleterious in our experiments, to simulate the distribution of (**A**) (five draws with replacement of the same number of elements as the real data). 10,000 simulations were run and the observed proportion (**B**, 60%) was never observed.

**Extended Data Fig. 5.**
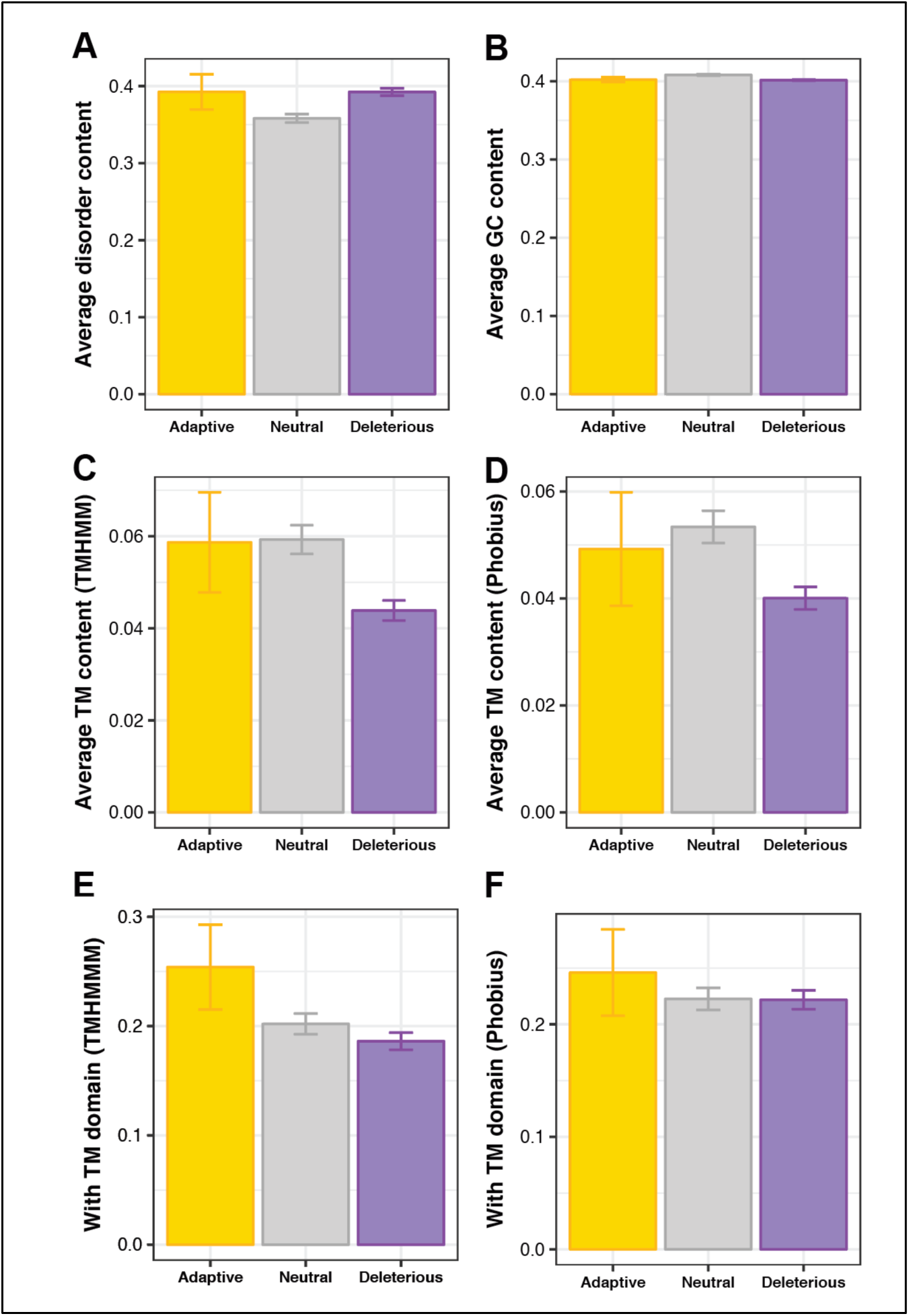
No association between TM propensity and beneficial fitness effects in established ORFs. The same analyses presented in Fig. 4 were repeated on the group of established ORFs. None of the 4 measures of TM propensity (C-F) are statistically different between adaptive and neutral established ORFs. The association between high TM propensity and beneficial fitness effects is specific to emerging ORFs (Fig. 4). See also Extended Data Fig. 6.

**Extended Data Fig. 6.**
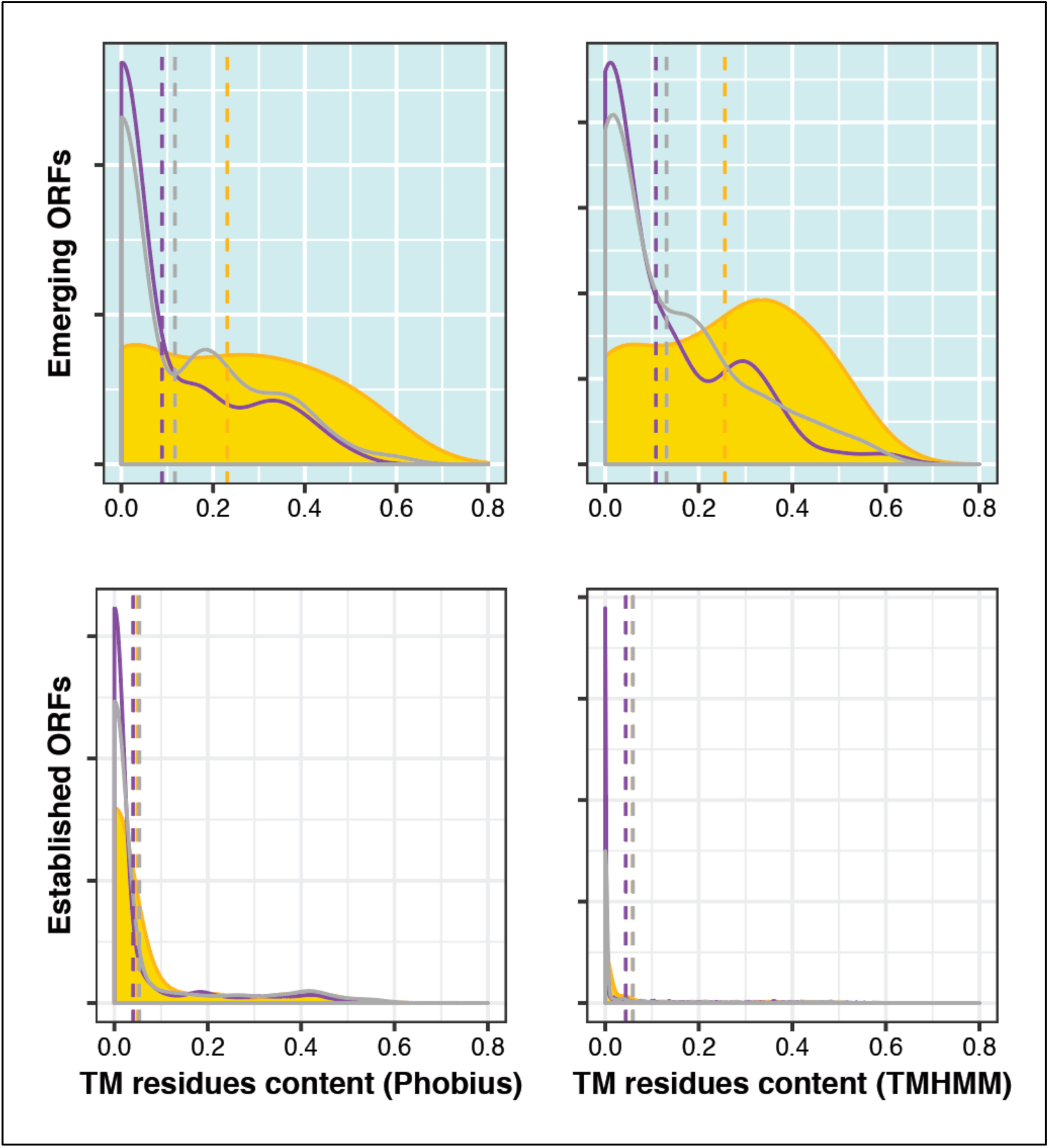
Distribution of TM residue content. Two prediction methods are compared: Phobius (left column) and TMHMM (right column). Two ORF classes are compared: emerging (top row) and established (bottom row) ORF. In each case, the distribution (density plot) of TM residue content (fraction of amino acids predicted as TM over length of the ORF) is shown for ORFs classified as adaptive (gold), deleterious (purple) and neutral (gray). Vertical dashed lines correspond to the mean of the distribution of the respective color.

**Extended Data Fig. 7.**
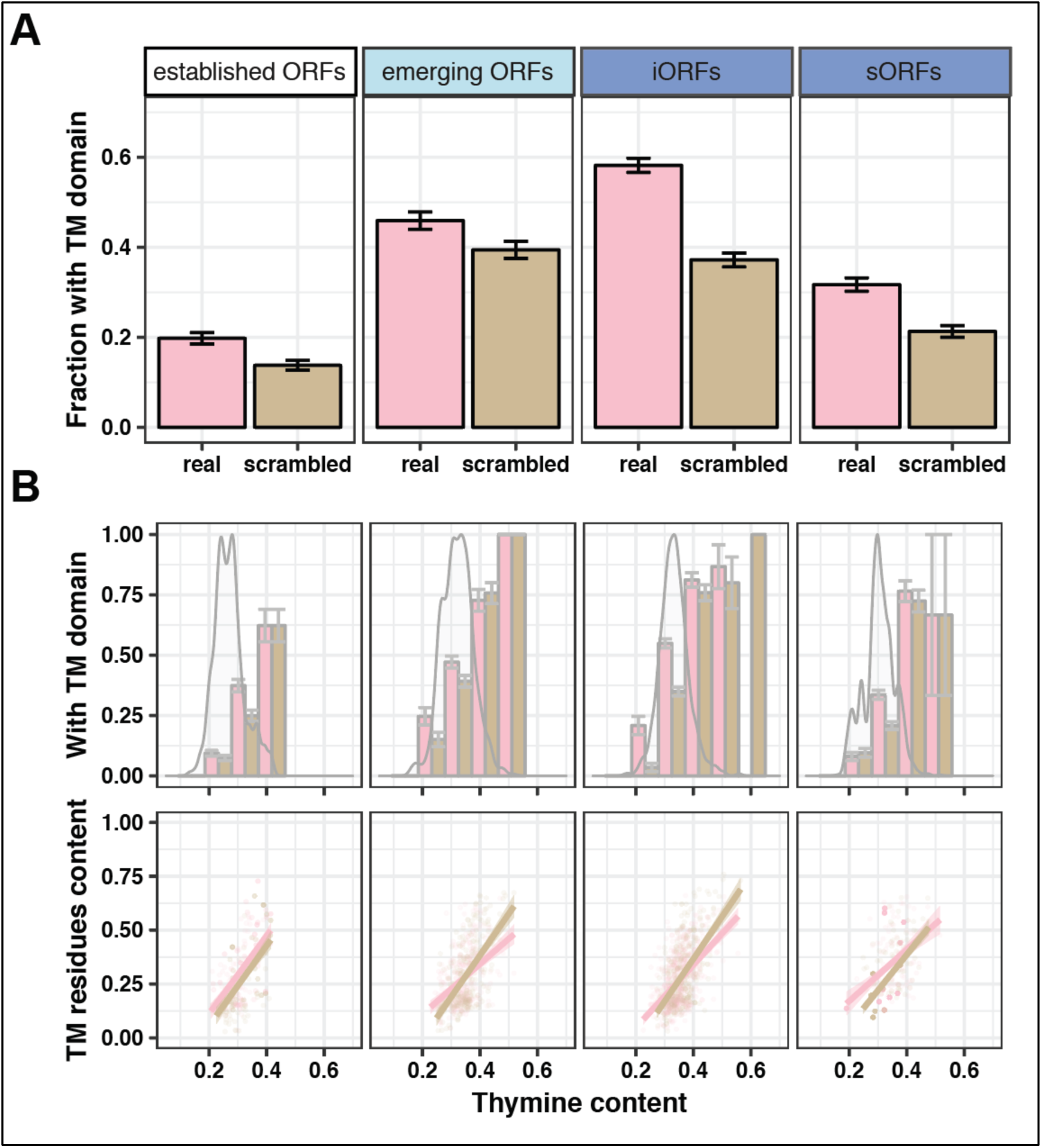
Analysis of TM propensity without sampling to control for length. **A.** <u>Propensity of ORFs to form TM domains.</u> Established ORFs, iORFs and sORFs have been sampled (n=1,000) with replacement to follow the length distribution of emerging ORFs. Error bars represent standard error of the proportion. **B.** <u>Thymine content influences TM propensity.</u> Top panel: Bar graph represents the fraction of sequences from (**A**) predicted to encode a putative TM domain, binned by thymine content (bin size = 0.1) and compared between real (pink) and scrambled (green) sequences. Error bars represent standard error of the proportion. Overlaid density plots represent the distribution of all sequences in each category. Bottom panel: scatterplot showing the fraction of sequence length predicted to be TM residues as a function of thymine content. Only sequences from (**A**) predicted to encode a TM domain are included in the bottom panel plots. Individual real (pink) and scrambled (brown) sequences are shown in the scatterplot with transparency; points of higher intensity indicate that sequences sampled multiple times. Linear fits with 95% confidence intervals are shown.

**Extended Data Fig. 8.**
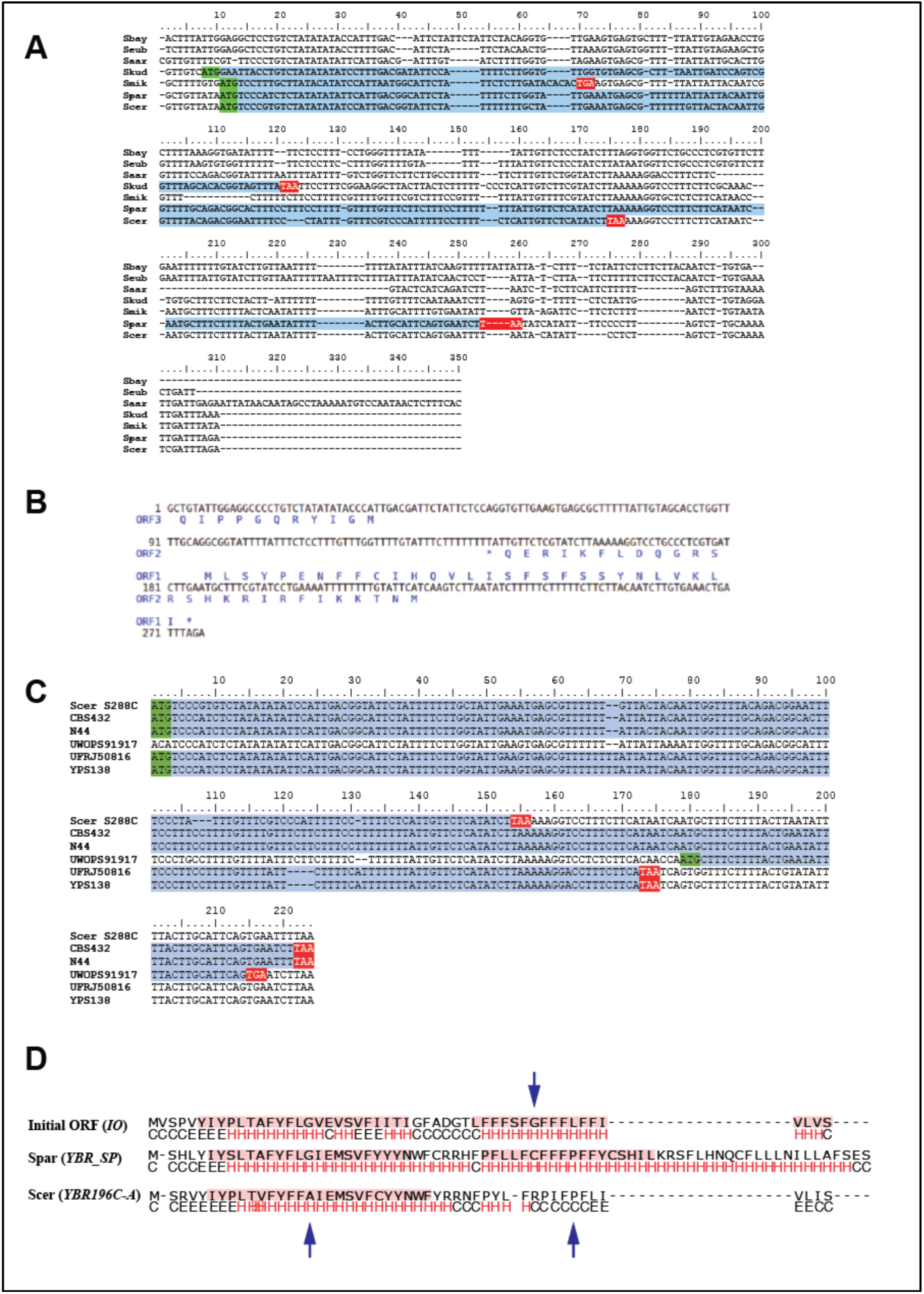
Extant and ancestral homologous regions to *YBR196C-A*. **A.** <u>Alignment of homologous regions to YBR196C-A in Saccharomyces species.</u> Start codons (green), stop codons (red), and ORFs (blue) are highlighted. Alignment was converted to Rich Text Format using BioEdit software (http://www.mbio.ncsu.edu/BioEdit/bioedit.html). **B.** <u>ORFs present at the reconstructed ancestral sequence.</u> Translation of ORFs (between a start and a stop codon) longer than 30 nucleotides is shown on the reconstructed ancestral sequence. ORFs identified using NCBI’s ORFfinder (https://www.ncbi.nlm.nih.gov/orffinder/). **C.** <u>Natural sequence variation in the S. paradoxus ortholog of YBR196C-A</u>. Start codons (green), stop codons (red), and ORFs (blue) are highlighted. Alignment was converted to Rich Text Format using BioEdit software and subsequently colorized. The S. cerevisiae reference sequence is aligned with the syntenic sequence in five S. paradoxus isolates. **D.** <u>Key sequence changes impair TM helices formation.</u> Alignment between translations of YBR196C-A (S. cerevisiae), YBR_SP (S. paradoxus) and YBR_IO (reconstructed initial ORF) with TMHMM predictions highlighted in pink and PsiPred secondary structure predictions shown underneath (H; helix; E: extended strand; C: coil). Arrows indicate helix breaking residues.

**Extended Data Fig. 9.**
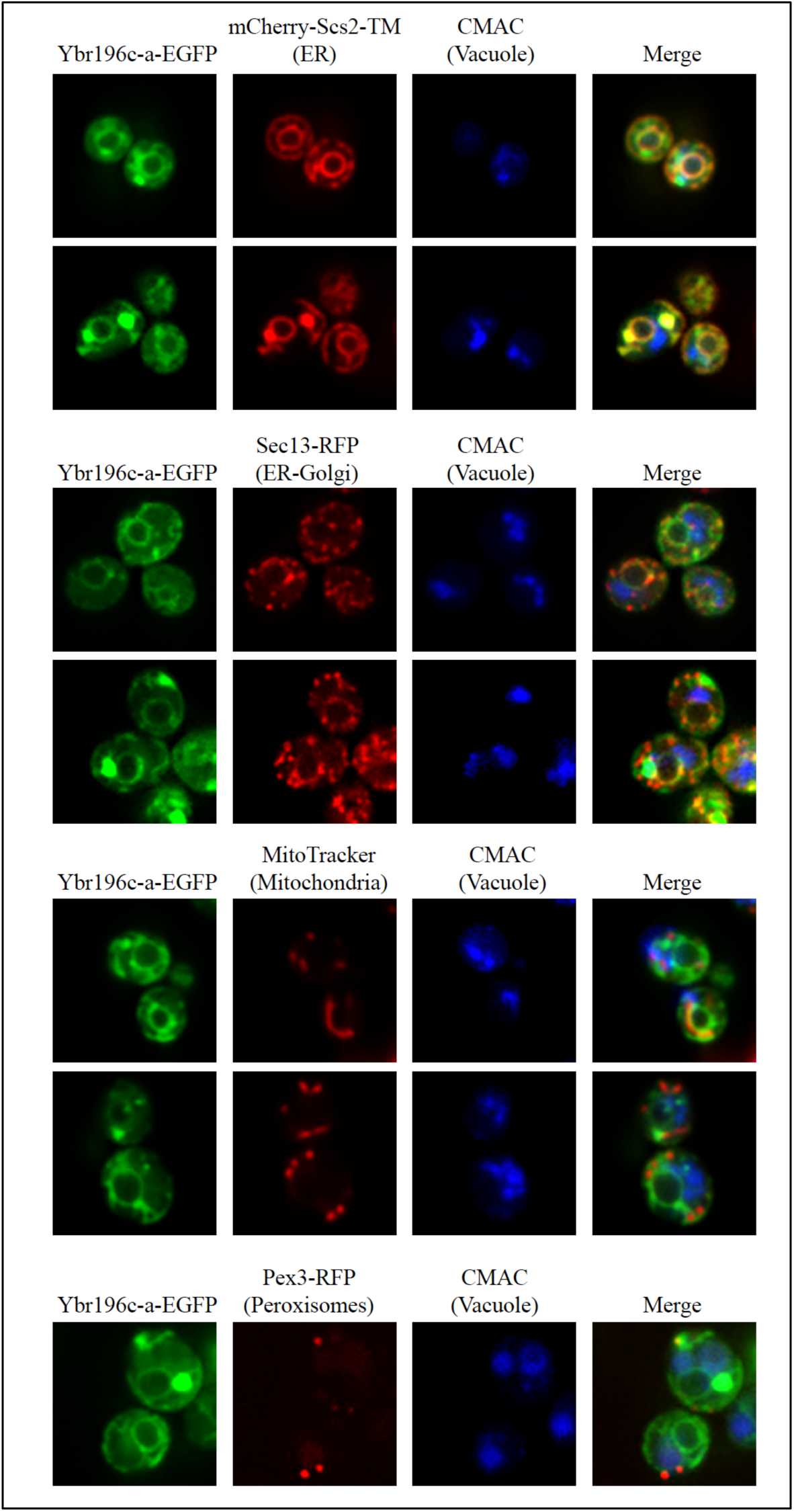
Co-localization studies with Ybr196c-a-EGFP. From top to bottom, plasmid-borne Ybr196c-a-EGFP and CMAC blue (to mark vacuoles) are visualized with markers of the ER (Scs2p), ER-Golgi (Sec13p), mitochondria (mitotracker) and peroxisomes (Pex3p). Images were acquired using confocal microscopy and representative micrographs are shown.

**Extended Data Fig. 10.**
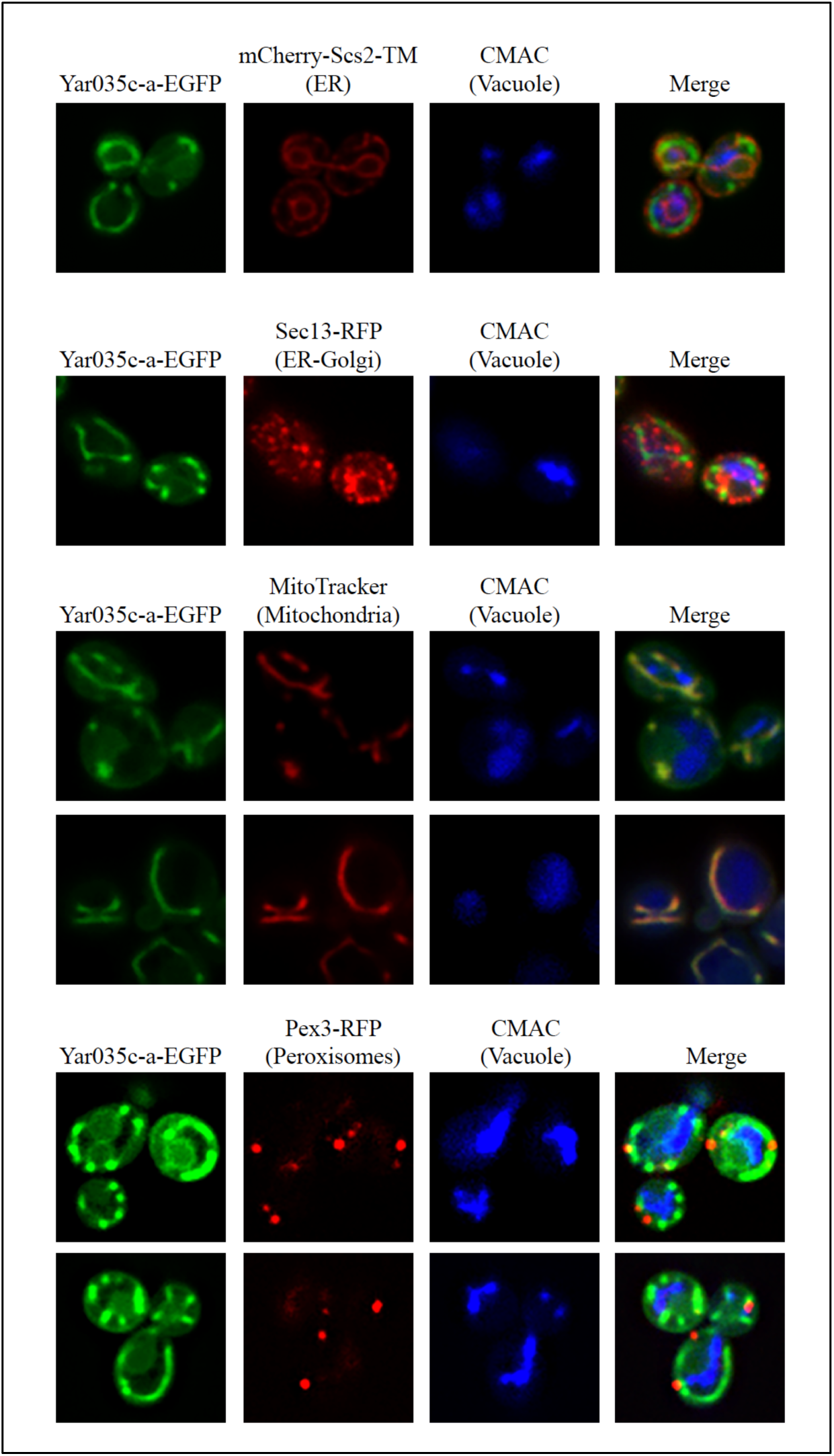
Co-localization studies with Yar035c-a-EGFP. From top to bottom, plasmid-borne Yar035c-a-EGFP and CMAC blue (to mark vacuoles) are visualized with markers of the ER (Scs2p), ER-Golgi (Sec13p), mitochondria (mitotracker) and peroxisomes (Pex3p). Images were acquired using confocal microscopy and representative micrographs are shown.

**Extended Data Table 1.** Yeast growth conditions.

| **Screening condition (N/C)** | **Complete name** | **Composition** |
| --- | --- | --- |
| N/A | SC+GLU+G418+5FOA | Synthetic complete + 2% Glucose + 100µg/mL G418 + 0.01% 5-FOA |
| N/A | SC+GLU+G418 | Synthetic complete + 2% Glucose + 100µg/mL G418 |
| N/A | SC+GAL+G418 | Synthetic complete + 2% Galactose + 100µg/mL G418 |
| N/A | SC-URA+GLU+G418 | Synthetic complete (-uracil) + 2% Glucose + 100µg/mL G418 |
| +/+ | SC-URA+GAL+G418 | Synthetic complete (-uracil) + 2% Galactose + 100µg/mL G418 |
| + /++ | SC-URA+GAL+RAF+G418 | Synthetic complete (-uracil) + 2% Galactose + 1% Raffinose + 100µg/mL G418 |
| ++/+ | SC-URA+CASE+GAL+G418 | Synthetic complete (-uracil) + casamino acids + 2% Galactose + 100µg/mL G418 |
| ++/++ | SC-URA+CASE+GAL+RAF+G418 | Synthetic complete (-uracil) + casamino acids + 2% Galactose + 1% Raffinose + 100µg/mL G418 |
| - /++ | SD-URA+GAL+RAF+G418 | Synthetic defined (+Histidine, Lysine, Leucine) + 2% Galactose + 1% Raffinose + 100µg/mL G418 |

SC (Synthetic complete) and SD (Synthetic defined) media are composed by 0.175% Yeast Nitrogen Base without amino acids and without ammonium sulfate and supplemented with 0.1% L-Glutamic Acid and 0.2% dropout amino acids. The dropout mixes were made with 10g of Leucine, 3g of Adenine, 0.2g of para-aminobenzoic acid and 2g of the remaining amino acids for each dropout mix.

**Extended Data Table 2.**
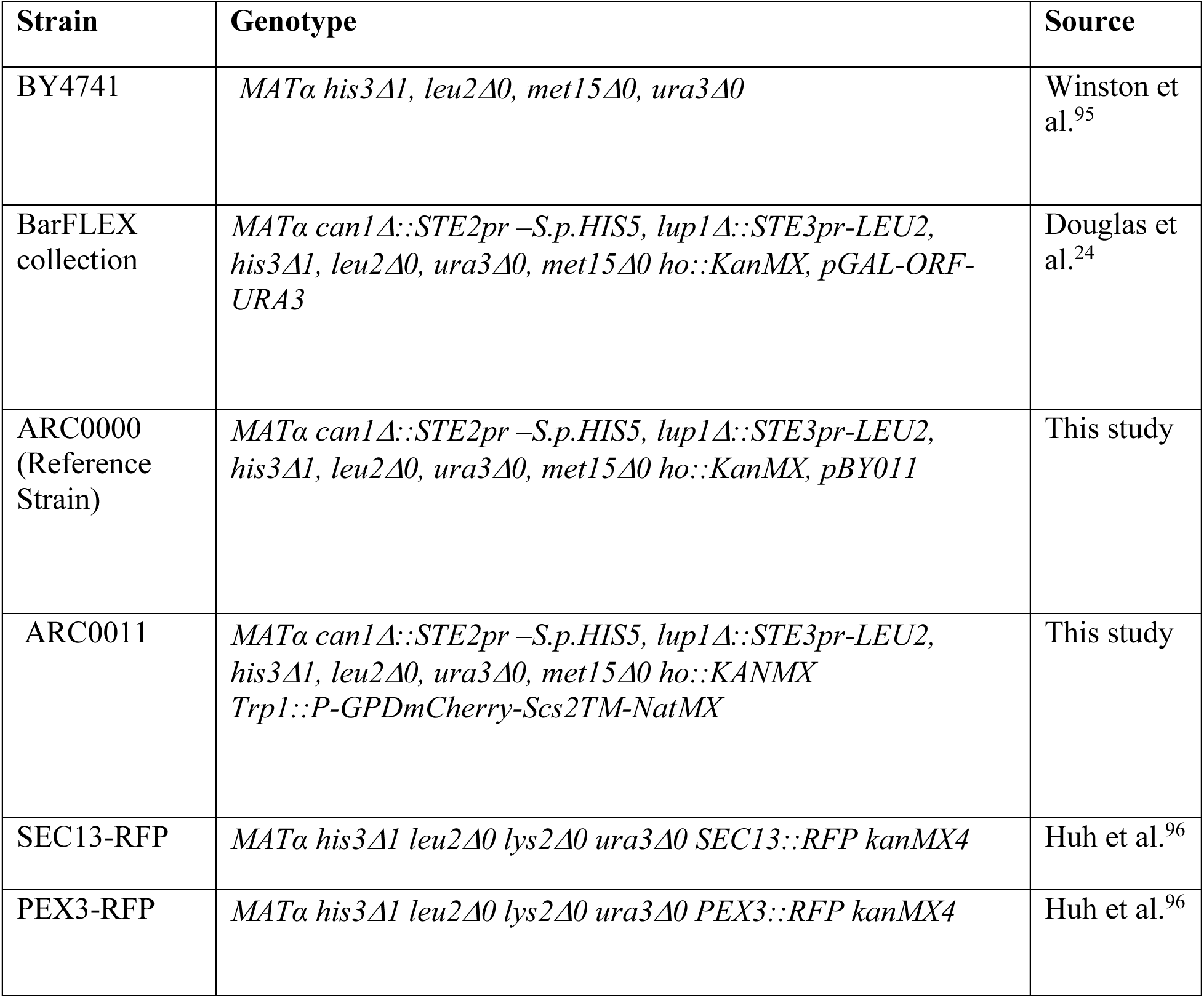
Genotypes of yeast strains used in this study.

**Extended Data Table 3.** Plasmids used in this study.

| **Plasmid** | **Description** | **Source** |
| --- | --- | --- |
| pBY011 | Yeast Gateway expression vector with ARS1, CEN4, Gal1-10 promoter (URA3) | Dana-Farber/Harvard Cancer Center |
| pDONR223 | Yeast Gateway donor vector | Rual et al.<sup>97</sup> |
| pDONR223-*YBR196C-A* | Entry Clone created by BP recombinase between the donor vector pDONR223 and the ORF *YBR196C-A* | This study |
| pDONR223-*YAR035C-A* | Entry Clone created by BP recombinase between the donor vector pDONR223 and the ORF *YAR035C-A* | This study |
| pAG426GAL-ccdB-EGFP | Yeast Gateway expression vector with 2μ, Gal1 promoter, EGFP (URA3) | Alberti et al.<sup>98</sup> |
| pAG426GAL-*YBR196C-A*-EGFP | Expression vector created by LR recombination between the destination vector pAG426GAL-ccdB-EGFP and Entry Clone pDONR223-*YBR196C-A* | This study |
| pAG426GAL-*YAR035C-A*-EGFP | Expression vector created by LR recombination between the destination vector pAG426GAL-ccdB-EGFP and Entry Clone pDONR223-*YAR035C-A* | This study |
| pSM3149 | p404 TRP P-GPDmCherry-Scs2TM-NatMX | Zhou et al.<sup>99</sup> |

